# A Prefrontal Cortex Map based on Single Neuron Activity

**DOI:** 10.1101/2024.11.06.622308

**Authors:** Pierre Le Merre, Katharina Heining, Marina Slashcheva, Felix Jung, Eleni Moysiadou, Nicolas Guyon, Ram Yahya, Hyunsoo Park, Fredrik Wernstal, Marie Carlén

## Abstract

The intrinsic organization underlying the central cognitive role of the prefrontal cortex (PFC) is poorly understood. The work to date has been dominated by cytoarchitecture as a canvas for studies on the PFC, constraining concepts, analyses, results, and their interpretations to pre-configured delimitations that might not be relevant to function. We approached organization by profiling the activity and spatial location of >23,000 neurons recorded in awake mice. Regularly firing neurons were over-represented in most PFC subregions, yet a fine-grained activity map of the PFC did not align with cytoarchitecturally defined subregions. Instead, we observed a robust relationship between spontaneous activity patterns and intra-PFC hierarchy, suggesting internal organization principles transcending cytoarchitecture. Single neuron responses to sounds did not reflect intra-PFC hierarchy but were linked to spontaneous firing rate, indicating that responsiveness increases with excitability and is decoupled from the PFC’s intrinsic operational structure. Our data-driven approach provides a scalable roadmap to explore functional organizations in diverse brain regions and species, opening avenues to link activity, structure, and function in the brain.

## Main

The prefrontal cortex (PFC) integrates brain-wide information and is crucial for emotional and cognitive functions^1,2^. While mapping of gene expression^3–5^, cytoarchitecture^6–8^, and connectivity^9–13^ provide insights to the structure of the PFC, a functional description of the PFC remains elusive^14^. It is unclear to which extent subregions of the PFC hold functional specializations^14,15^, and the existing structural descriptions of the PFC are yet to be integrated with the neuronal activities underlying this region’s information processing^16^. Furthermore, it is an open question whether the PFC’s neuronal activity is distinctive from other cortical regions and what functional demands PFC-specific activity patterns would support.

In the current study, we approach the organization of the PFC at the level of single neuron activity. A neuron’s firing pattern is a reflection of intrinsic biophysical properties and the neuron’s embedding in a particular network^17–19^. The activity profile of neurons should therefore inform about the functional properties of brain regions and could even improve their delineation. In line with this, it has been shown that brain regions differ in activity patterns^18,20,21^ and that there is a correlation between anatomical hierarchy and firing patterns^22–24^. Yet, would brain (sub)regions as defined by cytoarchitecture – and hitherto used to spatially sample and summarize activity – match with a functional map of the brain? In the case of the PFC, whose functions are considered to be distributed rather than localized^25^, the question of whether there exists an internal functional organization in space – and, if present, whether this organization corresponds with cytoarchitecturally defined subregions – is particularly debated.

Here, we recorded single neuron activity using high-density probes (Neuropixels) in awake mice and evaluated neuronal firing patterns at multiple spatial scales. To account for potential differences between intrinsically and externally driven states^22^, we analyzed spontaneous activity and sensory-evoked responses separately. We address the general relationship between single unit activity and brain anatomy, as well as the PFC’s distinctive activity characteristics and internal structure. The approach presented is a roadmap, applicable across brain regions and species.

### Capturing the diversity of spontaneous firing patterns

We recorded the firing activity of 23,386 single units from awake, head-fixed mice (dataset KI, Fig. 1a–c). About half of the units originated from the PFC and half from other brain regions (3 cortical, 10 subcortical; Fig. 1d and Supplementary Data Fig. 1a,b). In PFC we included 11 subregions^9,14^: secondary motor area (MOs), anterior cingulate area – dorsal and ventral part (ACAd, ACAv), prelimbic area (PL), infralimbic area (ILA), orbital area – medial, lateral, and ventrolateral part (ORBm, ORBvl, ORBl), agranular insular area – dorsal and ventral part (AId, AIv), and frontal pole (FRP). As spontaneous activity, we extracted three-second (s) epochs preceding the onset of auditory stimuli (Fig. 1c, ∼227 epochs/recording session). Epochs contaminated by ‘sleep-like’ periods^26^ were excluded due to their fundamentally different, non-stationary firing characteristics (Supplementary Data Fig. 1c–e, Methods). While generally stationary within individual units, activity patterns varied considerably across units (Fig. 1d). To capture this diversity, we characterized each unit by three metrics: firing rate, burstiness, and memory. Burstiness reflects the variability of a unit’s inter-spike intervals, while memory is defined as the correlation coefficient between subsequent inter-spike intervals (Fig. 1e)^27^. Note that burstiness solely describes the distribution of inter-spike interval durations, while memory reflects the temporal ordering of inter-spike intervals. Memory and burstiness are mathematically independent from each other and together comprehensively capture two key aspects of firing patterns: sequential structure and regularity. These two metrics thus offer a robust characterization that encompasses the information provided by other commonly used interval metrics, such as CV2 and LvR^20^. The memory metric is related to neuronal timescales that recent studies^22,23^ derived from firing autocorrelations. However, the complexity of the autocorrelations at the single neuron level disagrees with parametrization into a timescale through exponential fitting^17^. In contrast, the memory metric used here avoids the assumptions underlying timescales and describes the sequential structure of firing in a less biased way. Each metric was computed per 3 s epoch and averaged across epochs, thus summarizing each unit’s firing pattern into three metrics (Fig. 1e,f and Supplementary Data Fig. 1f–i).

**Fig. 1.**
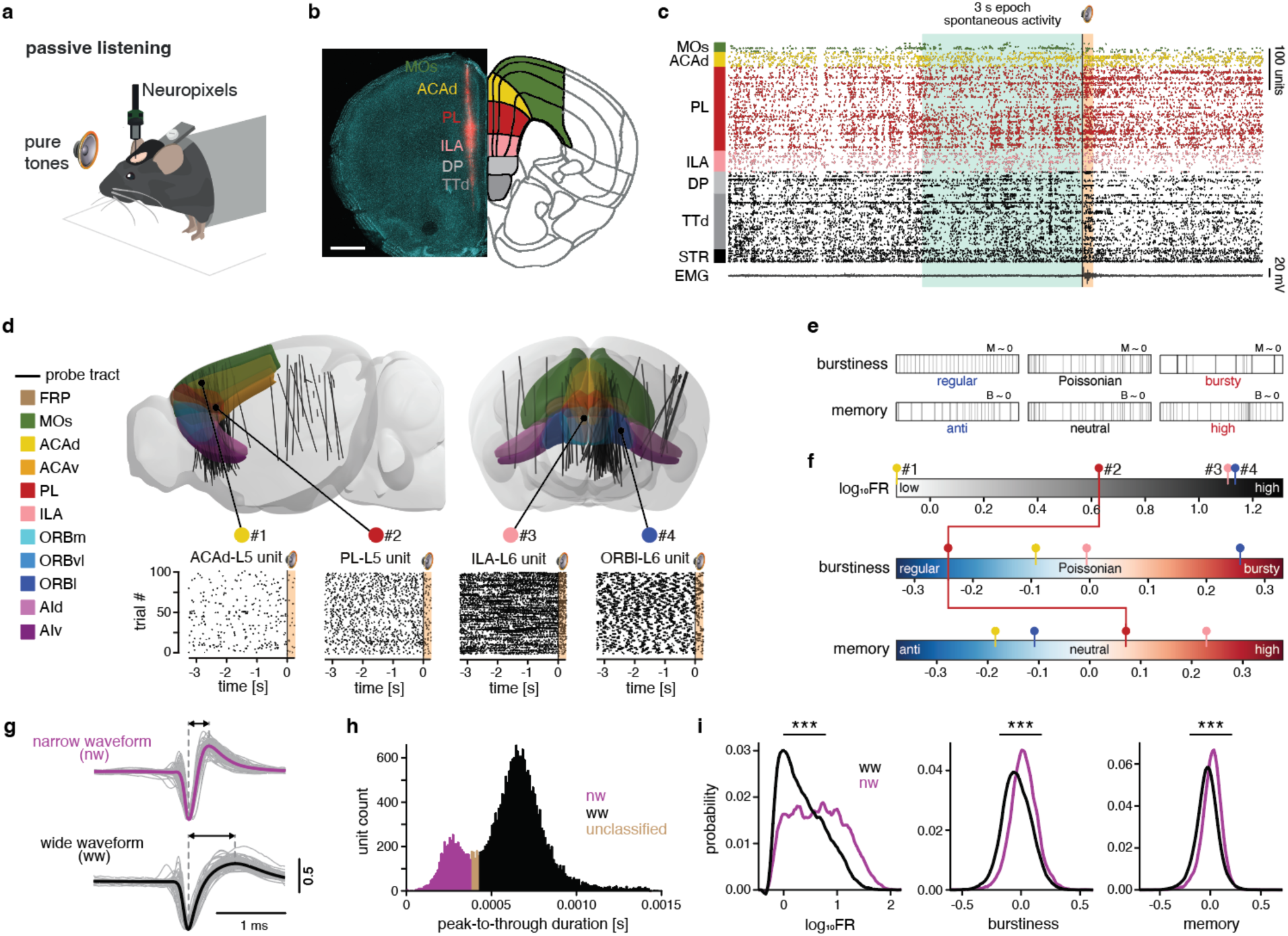
Characterization of spontaneous single unit firing patterns in the mouse brain. **a,** Experimental design; acute Neuropixels recordings in awake, head-fixed mice. Pure tones were presented during the sessions. **b,** *Left*, Coronal brain section with a Neuropixels probe track labeled by CM-DiI (red). Nuclei counterstained with DAPI (cyan). Scale bar 1 mm. *Right,* Corresponding plate in the Allen mouse brain reference atlas (CCFv3, plate #576989109) with color-coding of the sampled brain regions. **c,** Raster plot (*top*) of the 617 single units recorded with the probe shown in **b**. *Bottom,* electromyogram (EMG). Gray shading: 3 s epoch of spontaneous activity; orange shading: tone presentation (200 ms); black vertical line: tone onset. **d,** *Top*, Anatomical 3D localization of all probe tracts (n = 99, black lines) in distinct subregions of the PFC (colored) and other brain regions. *Bottom*, example spike raster plots (100 consecutive 3 s spontaneous epochs) of four single units (#1–4), recorded in layer 5 and 6 (L5, L6). **e,** Schematic illustrating the three firing metrics used to characterize spontaneous firing patterns. Each epoch (box) contains the same number of action potentials (i.e. same firing rate; 30 vertical bars). *Top row*, three cases of distinct burstiness: regular, Poissonian (random) and bursty with a constant neutral memory (M∼0). *Bottom row*, three cases of distinct memory: anti/neutral/high memory with a constant burstiness (B∼0). Adapted from ^27^. **f,** The three firing metrics used to characterize spontaneous activity of all recorded units: firing rate (log10FR *top)*, burstiness (*middle*), and memory (*bottom*). The gradient bars outline the respective metric range. The mean metrics across spontaneous epochs of the four units (#1–4) in **d** are indicated by colored circles and lines; the metric combination of unit #2 (red) is highlighted. **g,** Examples of individual (n = 100, grey) and mean (black/purple) waveforms of a narrow (nw, *top*) and a wide waveform (ww, *bottom*) unit. Double arrows indicate peak-to-trough durations. **h,** Distribution of peak-to-through durations of all mean waveforms (n = 23,667 single units; bin size: 1 ms). Peak-to-through duration < 0.38 ms = nw, n = 4200 units; > 0.43 ms = ww, n = 19,186 units. Units with intermediate peak-to-through duration were not classified (n = 281 units) and excluded from further analysis. **i,** Distributions of log10FR, burstiness, and memory for nw (purple) and ww (black) units. *** *p* < 0.001, mixed-effect regression (Methods).

Spike waveforms have been widely used to distinguish two neuron types, wide-width (ww) and narrow-width (nw) units (Fig. 1g), identified as putative excitatory and putative inhibitory neurons, respectively^28^. As expected, spike widths were bimodally distributed, allowing us to distinguish ww (n = 19,186; 81.1%) and nw (n = 4200; 17.7%) units (Fig. 1h). Nw neurons are expected to fire at high rates^28,29^, and indeed, our nw units displayed significantly higher firing rate, along with higher burstiness, and memory compared to ww units (Fig. 1i). To adequately analyze differences within each unit type, we processed ww and nw units separately. In the following, we focus on ww units.

### Brain regions have distinct firing repertoires

To obtain an overview of naturally occurring firing patterns, we trained a self-organizing map (SOM) on the three metrics extracted from the ww units. A SOM consists of a grid of nodes (in the current study hexagons), where each node represents a combination of metrics; similar nodes are neighbors on the map (Supplementary Data Fig. 2a). Our SOM displayed perpendicular gradients of burstiness and firing rate, indicating that these two metrics vary independently of each other. In contrast, high memory patches coincided with medium to low burstiness, indicating that high memory and high burstiness are mutually exclusive (Fig. 2a). Thus, burstiness and memory were empirically dependent, albeit mathematically independent. Note that linear correlation failed to reveal this empirical dependence between burstiness and memory (Supplementary Data Fig. 1f,g).

**Fig. 2.**
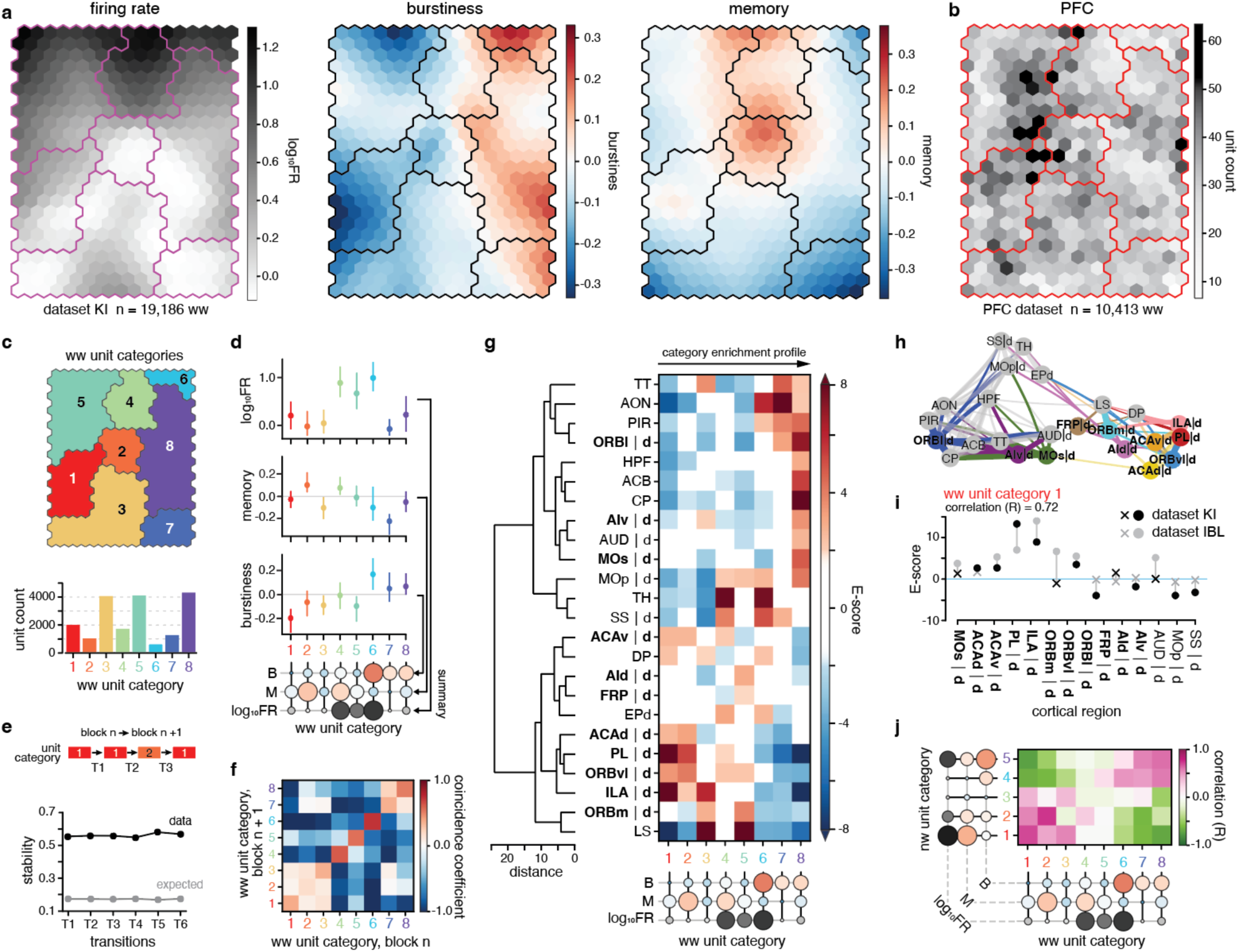
Brain regions have distinct firing patterns, with a low-rate, regular-firing signature for PFC subregions. **a,** The SOM’s component planes for the three metrics. Each component plane consists of a hexagonal grid of nodes and displays the respective metric value per node in color. Together, the component planes visualize the combined feature landscape of the SOM. Contours (black/purple) delineate the unit categories defined in **c**. **b,** Number of PFC ww units assigned to each SOM node. Contours (red) delineate the unit categories defined in **c**. **c,** *Top*, Partitioning of the SOM nodes into eight unit categories using hierarchical clustering. *Bottom*, number of ww units per unit category. **d,** Summarizing the characteristics of each unit category. Median (dot) and 10th to 90th percentile (vertical line) of metrics across units assigned to each category. Circles below (‘summary’) further summarize the metric composition of each category (filling: median metric value colored according to **a**, radius: relative magnitude of metric value compared to the other categories). **e,** Stability of ww unit categories across time. *Top,* Spontaneous epochs were split into blocks (∼50 epochs/block) and each unit’s category was calculated per block. *Bottom,* Stability (fraction of units retaining their category) from one block to the next (black) compared to stability expected from marginal distributions (grey). **f,** Quantification of the transitions between ww unit categories across all blocks shown as a coincidence coefficient matrix: -1: zero transitions; 0: random; 1: maximal possible number of transitions as derived from the marginal distributions (Methods). **g,** *Right*, Category enrichment profiles (Methods, Supplementary Data Fig. 3c) of brain (sub)regions. Bold: PFC subregions. |d : deep layers. Non-significant E-scores are whitened (Table 5). *Left*, hierarchical tree derived from the enrichment profiles. **h,** Graph representation of the data in **g**. Nodes representing brain (sub)regions are arranged according to the first and second UMAP dimension of their enrichment profiles, line width scales with cosine similarity between category enrichment profiles of regions (only shown for similarities > 0.1). Bold: PFC subregions. |d : deep layers. **i,** Comparison of the enrichment of ww category 1 units in cortical (sub)regions between our dataset (dataset KI, black) and a dataset from the International Brain Laboratory (dataset IBL, gray). Dots and crosses indicate significant and non-significant enrichment, respectively. Bold: PFC subregions. |d : deep layers. Correlation between datasets (KI and IBL): R = 0.72. **j,** Comparisons of ww and nw enrichment profiles. Color displays the Pearson correlation coefficient between enrichment in a ww and a nw category across the brain regions shown in **g**. *Data:* dataset KI, ww units, all brain (sub)regions and layers, n = 19,186 units; **a**–**f** dataset KI, ww units, all brain (sub)regions, for cortex restricted to deep layers (L5–6), n = 18,056 units; **g**, **h** dataset KI: ww units, cortical (sub)regions, deep layers (L5–6), n = 9,715 units; dataset IBL: ww units, cortical (sub)regions, deep layers (L5–6), n = 5,783 units; **i** dataset KI, nw units, all brain (sub)regions, for cortex restricted to deep layers (L5–6), n = 3,984 units; **j**

Each unit was assigned to the node of the SOM that matched its metrics best. Each node represented units from numerous recordings, confirming that inter-animal variability did not drive the SOM’s feature landscape (Supplementary Data Fig. 2b). While PFC units constituted 54% of the ww dataset used to create the SOM, units from the other brain regions were represented similarly well by the SOM (Supplementary Data Fig. 2c,d). Overall, PFC units were widely distributed on the SOM, with many units matching best to regularly-firing (i.e. low-burstiness) and avoiding anti-memory areas (Fig. 2b), while units from the other brain regions populated different areas (Supplementary Data Fig. 2e). Notably, occupancy of SOM areas varied across PFC subregions, indicating spatial differentiation of firing repertoires within the PFC (Supplementary Data Fig. 2f).

### Stable classification of single unit firing patterns

To enable a statistical analysis of the spatial distribution of firing patterns, we categorized firing patterns by hierarchically clustering the SOM’s nodes – and, implicitly, the matched units (Fig. 2c and Supplementary Data Fig. 3a,b). Units within a category shared similar firing statistics (Fig. 2d). To test whether units maintained stable firing properties over time, we allocated the spontaneous epochs into equally sized blocks and determined each unit’s category in the respective block. The category labels (1–8) assigned to a unit per block matched well with the unit’s category obtained across all epochs (Supplementary Data Fig. 3c). Consistently, around 57% of units remained in the category they had in the previous block, which is considerably above the 18% expected from chance (marginal distributions; Fig. 2e). Splitting data into blocks reduces the sampling of spikes, deteriorating metric estimation and consequently also categorization (clustering) of units. This should disproportionally affect low-rate units, leading to underestimation of the stability for units with low firing rate in particular. Indeed, low-rate categories (1–3, 7, and 8) were only about half as stable as the high-rate categories (Fig. 2f and Supplementary Data Fig. 3d). The low-rate, regular-firing categories (1–3) mostly transitioned among themselves, as did the low-rate, bursty categories (7 and 8), suggesting the existence of statistically similar category groups.

### Enrichment of low-rate, regular-firing units in the PFC

To quantify differences in firing patterns across various spatial entities – brain (sub)regions, layers, regions-of-interest (ROIs) – we defined a variable, the E-score, expressing for any selection of units the over/under-representation of a given unit category (1–8). More specifically, the E-score quantifies how many units belong to a given category (1–8) relative to a chance distribution. The magnitude of a positive/negative E-score reflects the statistical significance of a category’s enrichment/depletion (Supplementary Data Fig. 3e). E-score analysis of ww unit activity revealed substantial enrichment in bursty units in the superficial cortical layers (L2/3) while deep cortical layers (L5–6) were enriched in regular-firing units (Supplementary Data Fig. 3f,g and Supplementary Data Tables 1 and 2). Given the anatomical organization of the mouse PFC and the experimental constraints, 95% of cortical units were recorded from deep layers. Therefore, the subsequent analyses include all units from non-cortical regions, along with only the deep layer units from cortical regions.

The majority of the cytoarchitectural PFC subregions, i.e., ILA, PL, ACAd, ACAv, ORBm, and ORBvl, were generally enriched in low-rate, regular-firing units (categories 1–3), while holding subregion specific differentiations (Fig. 2g). Interestingly, ILA and PL were both heavily and significantly enriched in units of the most regular-firing category 1 (Supplementary Data Table 3). The ORBl, MOs, and AIv, in contrast, were significantly enriched in bursty category 8 units. Thus, the subregions ILA, PL, ACAd/v, and ORBm/vl, formed an activity-defined prefrontal entity, excluding ORBl, MOs, and AIv (Fig. 2h). Other brain regions like thalamus (TH) and hippocampal formation (HPF) were heavily and significantly enriched in bursty units (categories 6–8), revealing a PFC-specific signature of low-rate, regular-firing units. Characterizing brain regions by firing properties requires reproducibility and robustness across datasets. Repeating the analyses on the dataset of the International Brain Laboratory (dataset IBL; n = 25,066 ww units, for cortex L5–6) revealed a high degree of similarity between the regional enrichment profiles of the two datasets (Fig. 2i and Supplementary Data Fig. 4).

To analyze nw unit activity, we trained a separate SOM, and then repeated categorizations and comparisons of brain regions (Supplementary Data Fig. 5). ILA, PL, and ACAd/v were significantly enriched in high-rate, regular-firing nw units with high memory (Supplementary Data Fig. 5g and Supplementary Data Table 4). ORBl was consistently and significantly enriched in high-rate, bursty units. Considering both nw and ww unit activity, we found regular firing patterns to be the hallmark of the medial wall PFC subregions (ILA, PL, and ACAd/v). Relating nw and ww category enrichment profiles, we observed that brain (sub)regions enriched in low-rate, regular-firing ww units were enriched in high-rate, regular-firing nw units. Similarly, regions enriched in low-rate, bursty ww units were enriched in high-rate, bursty nw units (Fig. 2j).

### Category enrichment reflects cortical hierarchy

Having found differences between brain regions in terms of firing patterns, we next asked whether these differences are linked to the hierarchical organization of brain regions. The Allen Mouse Brain Connectivity Atlas describes the hierarchy of mouse cortical regions based on input/output connectivity motifs occurring between cortical and thalamic regions^9^. To investigate the relationship between our spontaneous activity categories (1–8) and cortical hierarchy, we correlated the E-scores of cortical (sub)regions with their respective anatomical hierarchy score established in this resource.

We found a positive correlation between anatomical hierarchy and enrichment in low-rate, regularly-firing units (categories 1–3). This positive correlation was not driven by a continuous relationship but due to a bimodal distribution of hierarchical scores between the low-hierarchy sensory regions versus the high-hierarchy PFC subregions (Fig. 3a and Supplementary Data Fig. 6a). To sample a wider range of cortical regions and hierarchical scores, we repeated the analysis with the dataset IBL. While individual cortical regions’ E-scores could differ between dataset KI and IBL, the positive correlations between hierarchy and the PFC’s signature categories 1–3 were corroborated (Fig. 3b and Supplementary Data Fig. 6b). We identified even stronger, negative correlations between anatomical hierarchy and enrichment in bursty, low-memory units (categories 6–8, Fig. 3c and Supplementary Data Fig. 6a,b). Importantly, when we limited the correlation analysis to PFC subregions, we found no direct relationship between hierarchy and enrichment/depletion in unit categories (Supplementary Data Table 5), suggesting that firing patterns did not reflect intra-PFC hierarchy at subregional level.

**Fig. 3.**
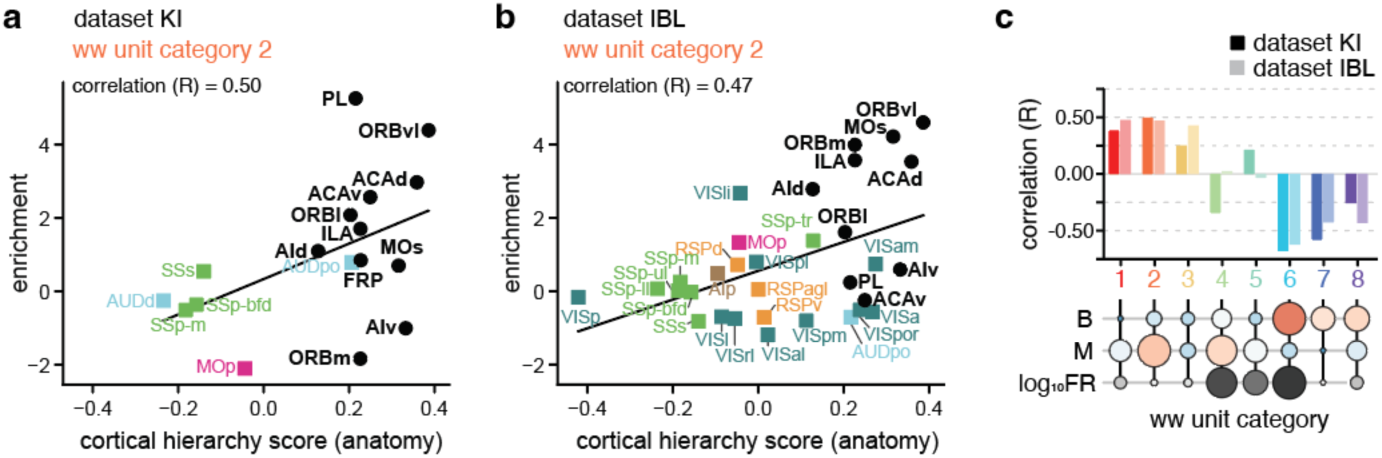
Spontaneous firing patterns reflect anatomical hierarchy of cortical regions. **a, b,** Enrichment in category 2 units vs cortical hierarchy score for our dataset (dataset KI, Pearson correlation: R = 0.50, **a**) and the dataset IBL (R = 0.47, **b**). Bold: PFC subregions. **c,** The Pearson correlation coefficient between cortical hierarchy and enrichment in the eight categories for our dataset (dataset KI, dark colors) and the dataset IBL (light colors). *Data:* dataset KI, ww units, cortical (sub)regions, deep layers (L5–6), n = 10,898 units; **a**, **c** dataset IBL, ww units, cortical (sub)regions, deep layers (L5–6), n = 7,168 units; **b**, **c** Cortical hierarchy scores from Harris et al.^9^.

### Towards an activity defined map of the PFC

Given the accumulating support for incongruence between cytoarchitecture and connectivity of the mouse PFC^14^, demarcating and analyzing activity profiles based on cytoarchitecture could be called into question. We therefore parcellated the PFC subregions into smaller regions of interest (dataROIs, n = 42), each containing a similar number of units (∼200; Fig. 4a,b and Supplementary Data Fig. 7a), and calculated E-scores (ww, deep layers) per ROI (Methods). Low-rate, regular-firing units (categories 1 and 3) were significantly enriched in ROIs located in ILA and PL, while low-rate, bursty units (categories 7 and 8) were enriched in ROIs located in the MOs and the ORBl (Fig. 4c and Supplementary Data Table 6). Clustering ROIs according to category enrichment profiles enabled us to define and spatially outline activity-based modules. (Fig. 4c and Supplementary Data Fig. 7b). This revealed a homogenous enrichment profile in the ILA, creating an ILA-specific activity module that bordered a module with similar firing patterns in the adjacent parts of PL and ACAd (Fig. 4d). Other activity modules stretched over cytoarchitectural boundaries and occurred in multiple places. Thus, several cytoarchitectural subregions lacked spatially homogenous activity signatures. The strong enrichment of low-rate, bursty units (categories 7 and 8) characterizing ORBl (Fig. 2g) could now, due to the improved spatial resolution, be pinpointed to a ROI in the most ventral portion of this subregion. The similarity in the activity patterns in ILA and the adjacent portions of PL is consistent with the proposition of a ventromedial subdivision of the PFC defined by input and output connectivity^12,14^. We could not, however, identify specific activity signatures for the proposed dorsomedial and ventrolateral subdivisions. A confound could be that our focus on deep layers primarily captures output activity while the connectivity-based subdivisions of the PFC also consider the superficial input layers. Conversely, our relatively coarse sampling of the extensive territory of the dorsomedial (MOs and ACAd/v) and the ventrolateral regions (ORBl and ORBvl) precludes discovery of small, homogeneous activity modules.

**Fig. 4.**
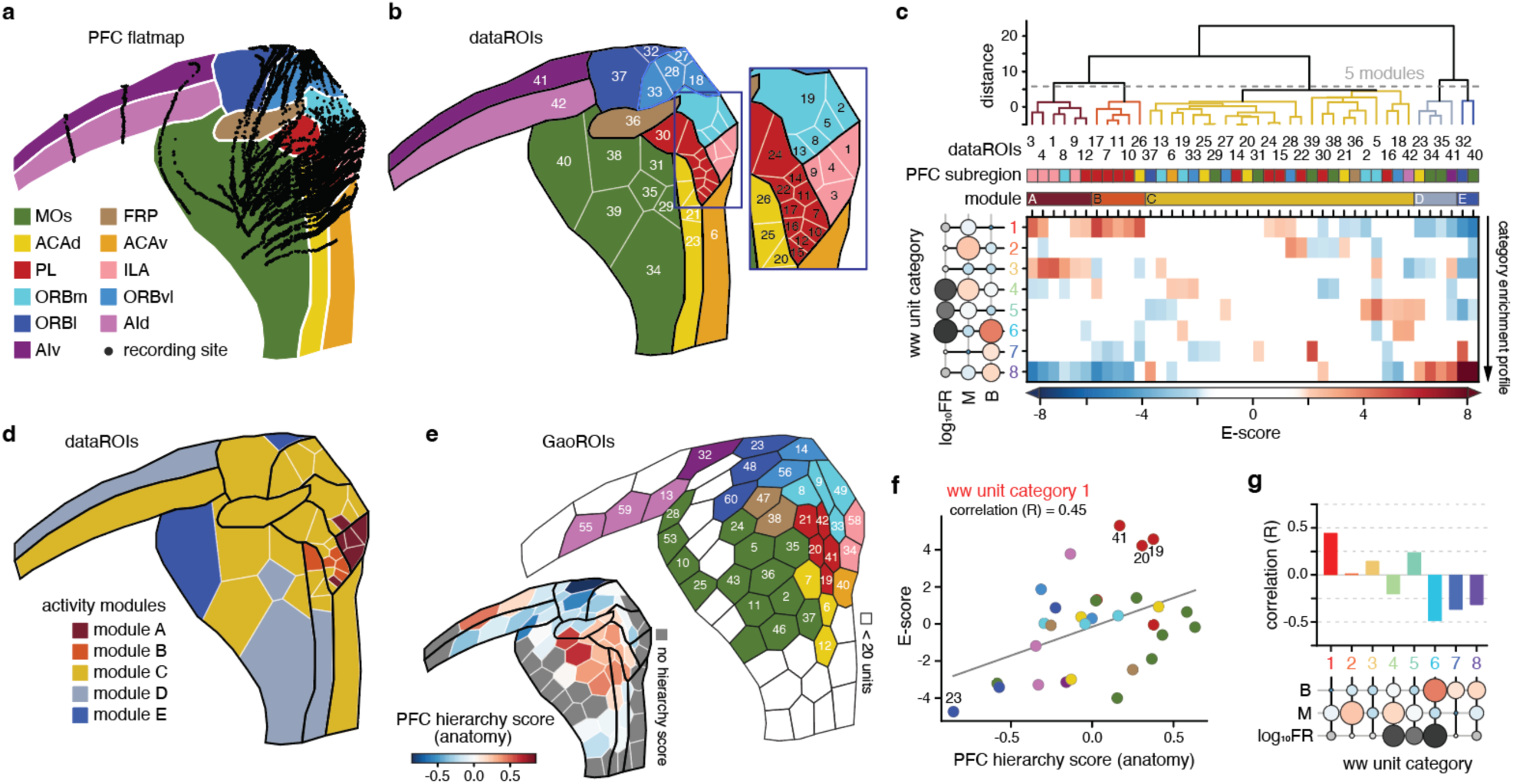
Activity-defined map of the PFC reflects intra-PFC hierarchy. **a,** Flatmap of the mouse PFC with the anatomical location of all recording sites (black dots, n = 99 Neuropixels probes) of the dataset KI. Colors: cytoarchitectural PFC subregions. **b,** PFC subregions (black outlines, colors as in **a**) parcellated into regions of interest (dataROIs, n = 42) holding similar unit counts (∼200 units per dataROI, Supplementary Data Fig. 7a). Each dataROI (white outlines) is identified by an ID number. Box: enlargement of the corresponding frame on the flatmap. **c,** Clustering of PFC dataROIs into activity modules based on their category enrichment profiles. *Top to bottom*, hierarchical tree derived from enrichment profiles; dataROI ID numbers; PFC subregion identity of the dataROIs (color-coded as in **b**); activity modules (A–E); category enrichment profiles of the PFC’s dataROIs. **d,** PFC flatmap with dataROIs (white outlines) colored according to activity module. Black outlines: cytoarchitectural PFC subregions. **e,** *Large flatmap,* the mouse PFC parcellated into regions of interest (GaoROIs, n = 60, black outlines) of roughly the same volume (mean ± s.d.: 0.288 ± 0.070 mm^3^). Each GaoROI is identified by an ID number and colored according to the PFC region where most of the ROI’s units were located (color see **a**). GaoROIs with fewer than 20 units were not considered (white). *Small flatmap*, GaoROIs colored according to PFC hierarchy score. **f,** Enrichment in category 1 units vs PFC hierarchy score (Pearson correlation: R = 0.44, P = 0.016). One dot = one GaoROI, colored as in **e**. **g,** Pearson correlation between PFC hierarchy score and category enrichment across GaoROIs. *Data:* dataset KI, PFC ww units, deep layers (L5–6), n = 9,319 units. PFC hierarchy scores in E to G from Gao et al.^10^.

Recently, two studies from Gao et al.^10,11^ used single neuron connectivity tracing to describe the mouse intra-PFC connectivity in detail and provided a map of the hierarchical organization of the PFC. Gao et al. parcellated the PFC into 60 ROIs of similar volume (GaoROIs) and found no apparent correspondence between hierarchy and cytoarchitecture. Using GaoROIs, the category enrichment profiles and activity-based modules obtained from dataROIs were replicated well (compare Fig. 4c,d with Supplementary Data Fig. 7c–f, and Supplementary Data Table 7). To determine how spontaneous firing patterns relate to the detailed intra-PFC hierarchy proposed by Gao et al., we correlated the E-scores of GaoROIs with their intra-PFC hierarchical score (Fig. 4e–g). Enrichment of low-rate, regular-firing units (category 1) was positively correlated to anatomical hierarchy (Fig. 4f) while enrichment in high-rate, bursty units (category 6) was negatively correlated to hierarchy (Supplementary Data Fig. 8). More specifically, GaoROIs (#19, 20, 41) located in PL displayed exceptionally high hierarchy scores and the strongest enrichment in category 1, while the GaoROI (#23) located in the ventral part of ORBl featured the lowest hierarchy score and most prominent depletion in category 1 (Fig. 4f). Unfortunately, no hierarchy scores were available for ROIs located in ILA, where we had observed a particularly strong enrichment in category 1 (Fig. 4c). Overall, our results demonstrate that the positive correlation between enrichment in regular-firing categories (1– 3) observed across cortical regions (Fig. 3a,c) is replicated within the PFC’s ROIs, indicating a general, scale-invariant link between enrichment in regular-firing units and hierarchy. Yet, as mentioned, the enrichment profiles of cytoarchitecturally defined PFC subregions had not shown an obvious relationship to anatomical hierarchy (Fig. 3a and Supplementary Data Fig. 6, and Supplementary Data Table 5). A partitioning scheme transcending the traditionally defined subregions might therefore better capture both activity and anatomical hierarchy in the PFC.

### No PFC specific signature in response to auditory stimulation

It is unclear how spontaneous activity translates to response activity at the single neuron level. Our recording sessions included presentation of 10 kHz tones (200 ms) separated by random inter-stimulus intervals of 5–10 s. From the dataset KI we obtained firing rate profiles of tone responses for all units with at least 20 response epochs (n = 15,352 units; Methods; Fig. 5a). In the following, we focus on ww units and restrict cortical regions to deep layers (L5–6). We normalized each response profile to the pre-stimulus period (Methods) and clustered the response traces according to their temporal profile into eight response categories (1–8), labeled according to the average peak delay. Response categories differed in response onset time, peak prominence, amplitude, and persistence (Fig. 5b,c and Supplementary Data Fig. 9a,b). Most units displayed flat or negative response traces (category 8). Aside from the flat/negative response cluster, we observed two distinct groups of response clusters. Response categories 1 to 4 formed a ‘homogeneous, high-amplitude’ group of small clusters, each displaying clearly localized peaks, consistent response peak onsets, and comprised of units with mostly significant responses. In contrast, response categories 5 to 7 formed a ‘heterogeneous, low amplitude’ group with variable peak onset latencies, persistent responses, and comprised of larger fractions of non-significantly responding units, each.

**Fig. 5.**
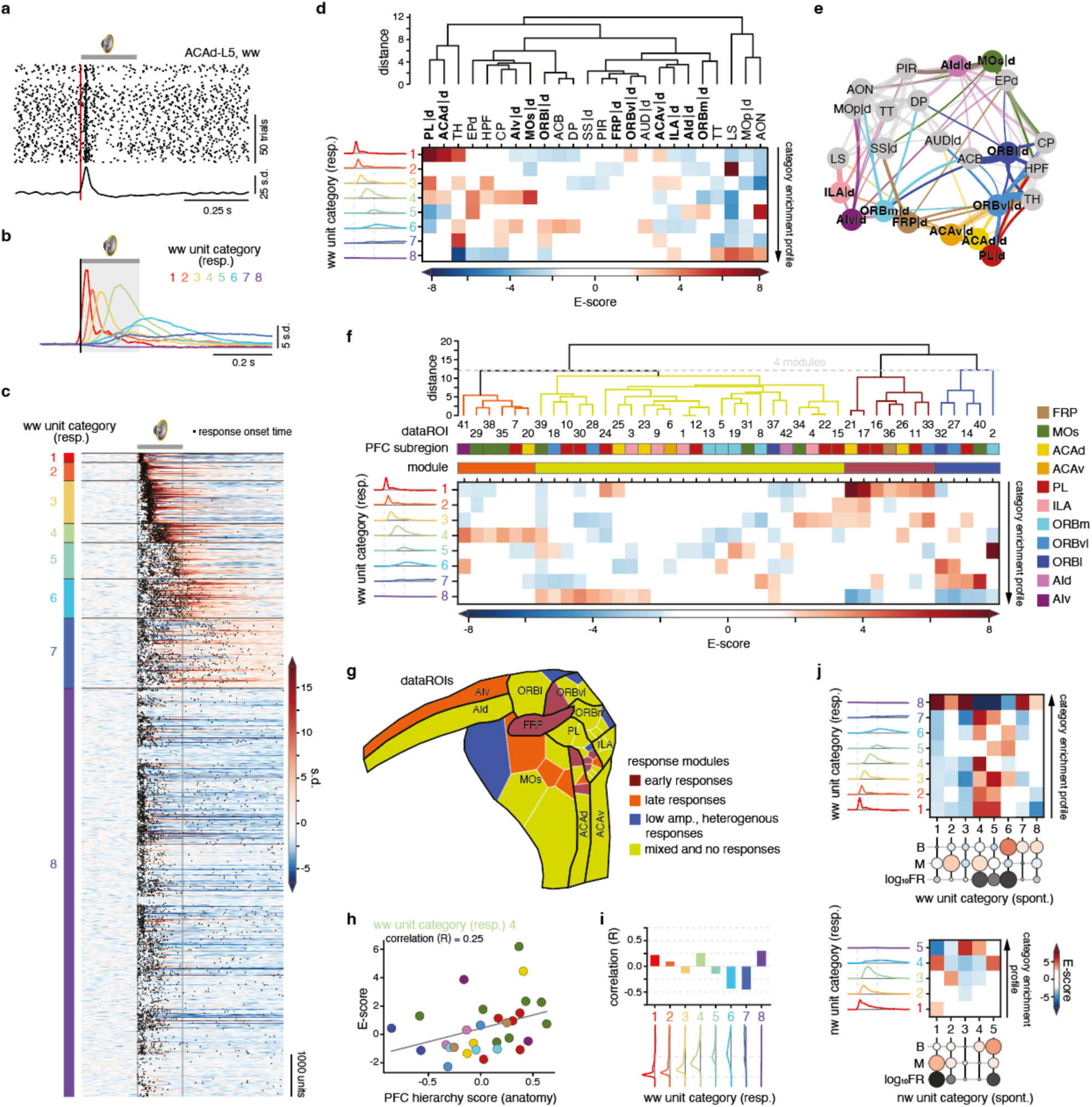
Single unit auditory responses in the PFC are constrained by spontaneous activity, rather than anatomy. **a,** Raster plot (*top*) of a layer 5 ACAd unit’s response to a 10 kHz tone (200 ms, gray horizontal line) and corresponding normalized peri-stimulus-time histogram (PSTH, *bottom*). Red vertical line: tone onset. **b,** The eight response categories of ww units. Each colored line represents the normalized PSTH averaged across the units assigned to a response category. **c,** Normalized PSTHs of single units arranged according to response category (1–8). Gray horizontal bar: tone presentation (200 ms); black vertical lines: tone onset/offset. Black dots: response onset time (ZETA test^39^, significant units only, P < 0.01). **d,** *Bottom,* Response category enrichment profiles of brain (sub)regions. Bold: PFC subregions. *Top,* hierarchical tree derived from the enrichment profiles. *Left,* average normalized PSTH per category as in **b**. Dashed vertical lines: tone onset/offset. **e,** Graph display of the data in **d**. Nodes representing brain (sub)regions are arranged according to the first and second UMAP dimension of their enrichment profiles, line width scales with cosine similarity between category enrichment profiles of brain regions (only shown for similarities > 0.1). Bold: PFC subregions. **f,** Clustering of PFC dataROIs (see Fig. 4b) into response modules based on their response category enrichment profiles. *Top to bottom*, hierarchical tree derived from dataROI enrichment profiles; dataROI ID numbers; PFC subregion identity of the dataROIs (colored as in Fig. 4b); response modules; response category enrichment profiles of the PFC’s dataROIs. **g,** PFC flatmap with dataROIs (white outlines) colored according to response module. Black outlines: cytoarchitecturally defined PFC subregions. **h,** Enrichment in response category 4 vs PFC hierarchy score (Pearson correlation: R = 0.25). One dot = one GaoROI, colored as in Fig. 4e. **i,** Pearson correlation between PFC hierarchy score and response category enrichment across GaoROIs. **j,** Relation between spontaneous activity patterns and auditory responses at the single-unit level. Matrices show how the eight different populations of spontaneous categories are differentially enriched in the eight response categories for ww units (top) and nw units (*bottom*). Enrichments are controlled for the influence of brain region (Methods, Supplementary Data Fig. 3c). *Data:* dataset KI, ww units, all brain (sub)regions and layers, n = 15,352 units; **a**–**c** dataset KI, ww units, all brain (sub)regions, for cortex restricted to deep layers (L5–6), n = 12,373 units; **d**, **e** dataset KI, PFC ww units, deep layers (L5–6), n = 7,184 units; **f**–**i**, **j** (*top)* dataset KI, PFC nw units, deep layers (L5–6), n = 1,486 units; **j** (*bottom)* PFC hierarchy scores from Gao et al.^10^.

To compare how units in different brain regions respond to auditory stimulation, we analyzed response category enrichment (E-score) profiles across cytoarchitecturally defined brain (sub)regions. In contrast to other brain regions (e.g., LS and AON), PFC subregions and thalamus were enriched in a variety of responsive unit types (response categories 1–7, Fig. 5d). PL, ACAd/v and ORBvl were significantly enriched in early responding units (response category 1), with PL and ACAd displaying the strongest enrichment. In addition, PL, ACAd/v, AIv, and ILA were enriched in the other homogenous, high-amplitude response categories 2 to 4. MOs showed a particularly strong enrichment in late, high-amplitude responding units (category 4). The PFC subregions ACAv, ORBm/l, AId/v were enriched in heterogenous, low-amplitude response categories 5 to 7. In summary, despite some commonalities, no response signature clearly demarcating and unifying the PFC across subregions emerged (Fig. 5e).

To generate a detailed response map of the PFC, we analyzed the response enrichment profiles of ROIs, applying the same approach used when previously generating the spontaneous activity map (Fig. 4b). Clustering the enrichment profiles of dataROIs into modules revealed strong within-subregion differences (Fig. 5f,g and Supplementary Data Fig. 9c). Large parts of the PFC were covered by a module reflecting enrichment in a mix of responding and non-responding units. PL’s and ACAd’s enrichment in early-responding units could now be attributed to a few early-responder ROIs within these subregions, which closely matched territories receiving direct input from AUD^12,13,30^. The most ventral parts of ORBl, ORBvl, and ORBm consistently featured heterogenous, low-amplitude responses (Fig. 5g) and shared a significant depletion in non-responding units (Fig. 5f). To relate the intra-PFC hierarchy to single unit responses, we correlated the response E-scores of GaoROIs with their intra-PFC hierarchical score. The anatomical hierarchy score showed a weak positive correlation with enrichment in early and late high-amplitude responding units (categories 1 and 4; Fig. 5h,i and Supplementary Data Fig. 9d–k). Conversely, the anatomical hierarchy score was strongly anti-correlated with enrichment in units pertaining to heterogenous, low-amplitude categories (categories 6 and 7). From previous work^22,23,31^, we expected to find high hierarchy scores, i.e. relatively more feedback connections, in ROIs and cortical (sub)regions enriched in units with sustained responses. In our data, units belonging to the high-amplitude category 4 and the low-amplitude heterogenous categories 5 to 7 could be interpreted as featuring sustained responses. Yet, these were differentially correlated with hierarchy, suggesting a multifaceted relation between hierarchy and response activity in the PFC.

### Spontaneous activity constrains response patterns

The maps derived from response and spontaneous activity strongly differed, raising questions about the direct relationship between spontaneous and response activity. We thus analyzed the response enrichment profiles of the eight spontaneous unit categories, statistically controlling for (sub)region influence (Fig. 5j and Methods). Low-rate units (spontaneous categories 1–3, 7, 8) consistently showed an exclusive enrichment in flat or negative responses (response category 8). Conversely, high-rate units (spontaneous categories 4–6) were enriched in various categories showing clear positive responses (response categories 1–7). These relationships also held true for nw neurons (Fig. 5j and Supplementary Data Fig. 10). This suggests that higher spontaneous firing rates reflect higher excitability and thus likelihood to respond.

## Discussion

In this study, we categorized spontaneous and response activity patterns of single units and evaluated enrichment of unit categories and correlation to anatomical hierarchy at two spatial levels. First, adhering to cytoarchitecturally defined brain regions, we addressed whether PFC subregions shared a unifying activity characteristic distinguishing the PFC from other brain regions. Second, we evaluated activity patterns in spatially fine-grained ROIs to challenge cytoarchitectural subregion boundaries within the PFC, yielding a first activity-based map of this region. Given the sampling constraints due to the anatomical positioning of the PFC, our analyses focused on layer 5 units when evaluating cortical regions.

We identified regular spontaneous activity as a common signature of the PFC’s subregions, with the exception of parts of ORBl and the MOs, which displayed enrichment in bursty units (Fig. 2i). Notably, the enrichment score reflects relative deviation from the sampled population. Hence, given the overrepresentation of PFC units in our dataset, we may have underestimated this signature’s prominence. Our findings differ from Mochizuki et al.^20^, who reported near-random (Poisson) firing in the PFC. This likely due to methodological differences, as Mochizuki et al. focused on in-task, non-stationary activity, analyzed fewer units in rodents, pooled units across layers and types, and used different delineations of brain regions. Regular firing patterns are functionally tied to cognitive processes, such as sustained input integration and stable representations (e.g., of context, rules and memories)^32,33^. Mechanistically, regular firing aligns with known PFC properties, including a high degree of recurrent connectivity and an abundance of ion channels with slow kinetics^5,34,35^. Future research should link spontaneous activity patterns to input and output connections, morphology, and genetic markers of individual units to deepen our understanding of these pattern’s origins and their functional impact.

Our observation that regions enriched in low-rate, regularly firing ww units were enriched in high-rate, regularly firing nw units (Fig. 2j) can be explained by differential targeting of ww and nw unit types by neuromodulatory inputs and long-range connections^36,37^. High-rate putative inhibitory nw units could also directly contribute to the low-rate characteristic of ww units, supporting the hypothesis that the PFC operates in an inhibition-dominated regime conducive to fine-tuned processing required for complex cognitive processes^37,38^.

The consistent correlations between anatomical hierarchy scores and enrichment in units of certain spontaneous activity categories across both the KI and the IBL dataset affirm a robust relationship between cortical hierarchy and spontaneous firing repertoire (Fig. 3). At the level of brain regions, this activity-hierarchy correlation is in line with studies on mice, macaques, and humans showing that spontaneous neuronal timescales correlate with connectivity-based anatomical hierarchy^23,24^. In contrast, Siegle et al.^22^ found no such correlation in the mouse visual system, potentially because (i) their timescale metric of single neuron activity did not reflect firing regularity – we show here that firing regularity, not memory, strongly drives correlations with anatomical hierarchy, (ii) the timescale metric was inadequate to capture the complexity of single unit autocorrelations^17^, and (iii) spontaneous activity-hierarchy correlations may be absent within the visual system.

In generating an activity-based PFC map, we observed that one spontaneous activity module aligned with the cytoarchitecturally defined ILA subregion, whereas other modules did not correspond to single subregions (Fig. 4d). Instead, spontaneous activity reflected connectivity-based anatomical hierarchy within ROIs (Fig. 4g), suggesting that connectivity, rather than cytoarchitecture, shapes the PFC’s activity landscape.

Unlike spontaneous activity, auditory response activity did not reveal a unifying PFC signature across subregions (Fig. 5d,e) and displayed a more multifaceted relationship to anatomical hierarchy. Our observation that the rate aspect of spontaneous activity predicts responsiveness (Fig. 5j), while the regularity aspect of spontaneous activity reflects spatial properties (Fig. 4), could explain the absence of spatial structure and regional signatures in response activity. Thus, spontaneous activity may better represent intra-PFC connectivity and spatially constrained biophysical properties, whereas auditory response activity rather reflects distributed, long-range connectivity^12,30^ from auditory regions.

The reproducibility of our results with the IBL dataset demonstrates the robustness of our approach across experimental settings and highlights the relevance of spontaneous firing patterns for elucidating brain organization. Our findings contribute to a better understanding of the PFC’s functional landscape, challenging the traditional reliance on cytoarchitecture by highlighting connectivity as a primary organizing principle of cortical activity. Our approach offers a framework for exploring other brain regions and species where cytoarchitectural and functional boundaries may similarly diverge.

## Methods

### Animals

All procedures and experiments on animals were performed according to the guidelines of the Stockholm Municipal Committee for animal experiments and the Karolinska Institutet in Sweden (approval numbers 7362-2019 and 1535-2024). 27 Male and 40 female adult wild type (C57BL/6J; Charles River) 3–6 months old mice were used. Animals were group housed, up to five per cage, in a temperature (23°C) and humidity (55%) controlled environment in standard cages on a 12:12 hours reversed light/dark cycle with *ad libitum* access to food and water, unless placed on a water restriction schedule. All water-restricted mice were restricted to 85–90% of their initial body weight by administering 1 mL of water per day.

### Surgery

Adult mice were anesthetized with isoflurane (3 % for induction then 1–2 %). Buprenorphine (0.1 mg/kg s.c.), carprofen (5 mg/Kg s.c.) and lidocaine (4 mg/kg s.c.) were administered. The body temperature was maintained at 37°C by a heating pad. An ocular ointment (Viscotears, Alcon) was applied over the eyes. The head of the mouse was fixed in a stereotaxic apparatus (Kopf). Lidocaine 2% was injected locally before skin incision. The skin overlying the cortex was removed, the skull was first cleaned with Chlorhexidine and then gently scratched with a scalpel blade. A thin layer of glue was applied on the exposed skull. A light-weight metal head-post was fixed with light curing dental adhesive (OptiBond FL, Kerr) and cement (Tetric EvoFlow, Invoclar Vivadent, Schaan, Liechtenstein). For extracellular recordings, a chamber was made by building a wall with dental cement along the coronal suture and the front of the skull. Brain regions were targeted using stereotaxic coordinates. After the surgery, the animal was returned to its home cage and carprofen (5 mg/kg s.c.) was provided for postoperative pain relief 24 hours following surgery.

### Habituation and behavioral settings

Following surgery recovery, mice were handled and progressively habituated to the head-fixation procedure over a period of 3–4 days by increasing the head-restriction time from 15 min to 1 h. After habituation, the mice were engaged in distinct behavioral settings. For each behavioral setting, auditory stimuli (10kHz pure tones) were delivered through earphones. Importantly, although the mice were engaged in various behavioral tasks, firing metric distributions were not significantly different across behavioral settings (linear mixed model). For all behavioral settings, delivery of the auditory stimuli was controlled with custom-written computer routines using a National Instruments board (PCI-6221) interfaced through Matlab (Mathworks).

### Head-fixed recordings

#### Preparation

For acute recordings, two to four small craniotomies (300–500 μm in diameter) were opened a few hours (> 3 h) before the experiment to access the pre-marked targeted entry points (PFC: +2.20 to +1.60 mm AP, ±0.3 to ±1.5 mm ML; AUD: -3.20 mm AP, ±4.20 mm ML; CA1: -1.50 mm AP, ±1.6 mm ML; MOp: +1.90 mm to +1.50 mm AP, ±2.50 mm ML; SSp: -3.10 mm AP, ±2.80 mm ML;). The mice were anesthetized with isoflurane (3 % for induction then 1–2 %). Buprenorphine (0.1 mg/kg s.c.), carprofen (5 mg/Kg s.c.) and lidocaine (4 mg/kg s.c.) were administered. The open craniotomy was covered with Silicone sealant (Kwik-Cast, WPI) and the mouse was returned to its home cage for recovery.

#### Probe preparation

The probes were coated with CM-DiI (1,1’-dioctadecyl-3,3,3’3’-tetramethylindocarbocyanine perchlorate, Thermo Fisher, USA) a fixable lipophilic dye for post-hoc recovery of the recording location. The coating was achieved by holding a drop of CM-DiI at the end of a micropipette and repeatedly painting the probe shank with the drop, letting it dry, after which the probe appeared pink.

#### Probe insertion

First, the electrode reference was then connected to a silver wire positioned over the pia in a craniotomy with a micromanipulator. Then, the probe(s) was(were) lowered gradually (speed ∼20 μ.s^-1^) with a micromanipulator (uMp-4, Sensapex, Oulu, Finland), using 0° to 20° angles, until the tip reached a depth of ∼3800–4200 μm under the surface of the pia. The probe(s) was(were) allowed to sit in the brain for 20–30 minutes before the recordings started. A total of 99 probes were lowered with a maximum of three probes inserted simulataneously.

#### Data acquisition

Extracellular potentials were recorded using Neuropixels probes (Phase 3B Option 1, IMEC, Leuven, Belgium) with 383 recording sites along a single shank covering 3800 μm in depth or with Neuronexus probes (A1x32-Poly2-10mm-50s-177) with 32 recording sites along a single 750 μm shank. The spike band data was digitized with a sampling frequency of 30 kHz with gain 500 while the LFP band data was digitized with a sampling frequency of 2.5kHz with gain 250. The digitized signal was transferred to our data acquisition system (a PXIe acquisition module PXI-Express chassis: PXIe-1071 and MXI-Express interface: PCIe-8381 and PXIe-8381, National Instruments for Neuropixels recordings or an Open Ephys acquisition board for Neuronexus probes), written to disk using SpikeGLX (Bill Karsh, Janelia) for Neuropixels probes or Open Ephys GUI for Neuronexus probes, and stored on local server for future analysis. The action potential signals (‘spike band’) were filtered between 0.3 Hz and 10 kHz and amplified. The local field potential signals (‘LFP band’) were filtered between 0.5Hz and 500 Hz.

#### Probe cleaning

After recordings probes were cleaned for 30 min with a fresh Tergazyme® solution (Sigma-Aldrich) and rinsed with distilled water overnight.

### Perfusion

At the end of the last recording session, each mouse was deeply anesthetized with pentobarbital (60 mg/kg; i.p.), and then transcardially perfused with 4 % paraformaldehyde (PFA). The brain was removed and post-fixed in 4% PFA in 1xPB.

### Tissue Clearing and Neuropixels probe track reconstruction

Brain sections were cut on a vibratome at 400 μm thickness (Leica VT1000, Leica Microsystems GmbH). Slices were repeatedly washed in PB and cleared using “CUBIC reagent 1 (25 wt% urea, 25 wt% *N,N,N’,N’*-tetrakis(2-hydroxypropyl) ethylenediamine and 15 wt% polyethylene glycol mono-*p*-isooctylphenyl ether/Triton X-100) for two days. After repeated washing in PB, slices were incubated with DAPI (1:50000) for one day at room temperature. Slices were then re-washed in PB and submerged in “CUBIC reagent 2” (50 wt% sucrose, 25 wt% urea, 10 wt% 2,20,20’-nitrilotriethanol and 0.1% v/v% Triton X-100) for further clearing. Slices were mounted on customized 400 μm thick slides using CUBIC2 solution and covered with 1.5 mm cover glasses. Blue and red channels were imaged at 4x magnification using a Zeiss 800 or 880 confocal microscope and exported via ZEN black (2.1 SP3 v14.0). For each brain section 6 to 7 z-stacks spaced by 50 μm were obtained and down-sampled to 10 μm resolution. The z-stacks containing the probe’s red fluorescent signal (DiI) were registered in the Allen Institute Common Coordinate Framework (CCFv3) and the probe position was estimated using the accompanying ‘SHARP-Track’ pipeline (https://github.com/cortex-lab/allenCCF). The electrode locations along the probe were transformed into CCFv3 space based on the orientation and position of the probe track. Unit locations were assigned based on the location of the electrode where that unit had the highest waveform amplitude.

### Spike sorting

#### Preprocessing

The high-pass filtered spike band data were preprocessed using common-average referencing: the channel’s median was subtracted to remove baseline offset fluctuations, then the median across channels was also subtracted from each channel to remove artifacts.

#### Kilosort2

The data was semi-automatically spike sorted with Kilosort 2.0 (https://github.com/MouseLand/Kilosort/releases/tag/v2.0). Clusters of waveforms were manually curated using the phy2 GUI (https://github.com/cortex-lab/phy). During manual curation, clusters of waveforms showing near-zero amplitudes, non-physiological waveforms, inconsistent waveform shapes and/or refractory period violations were discarded. The remaining units were classified as ‘good’. These potential ‘good’ units were finally compared with spatially neighboring clusters. Units showing similar waveforms, clear common refractory periods and putative drift patterns were subjected to a merge attempt: If the resulting cluster was still showing consistent waveforms and a clear refractory period in their auto correlogram, the merge was kept. Rarely, splits of clusters were performed when the principal features of waveforms indicated distinct clusters, and two or more groups of waveforms could be clearly identified. Double counted spikes were thus removed. Only the ‘good’ units were kept for the following analysis.

#### Spike-sorting quality control

pike times were corrected for temporal drifting during the recording time (∼10 ms/h) relative to a clock signal registered independently by the PXIe acquisition module and the PCI-6221 card logging the behavioral signals. First the temporal drift between the two devices was measured for each recording. Second, a linear regression was applied to correct the timestamps (behavioral timestamps) relative to the spike times. Spikes around saturation periods were detected with a margin of 1 s before and after saturation. Then, all units classified as ‘good’ were saved in Neurodata Without Borders (NWB) files for subsequent analysis.

#### Electromyogram (EMG) extraction

In Neuropixels recordings, we defined the EMG from the high frequency (1–10 kHz) muscular tone (neck muscles), picked up by our reference electrodes (e.g., over visual cortex) with zero time-lag^29,40,41^. EMG signals were extracted as the band pass filtered (ellipsoid band-pass filter with a lower and upper cutoff frequency of 1 kHz and 10 kHz respectively) median signal across all active channels (common noise source). EMG signals were down-sampled to 1 kHz and stored in NWB files for further analysis.

### Data analysis

Unless stated otherwise, continuous variables are reported as the median and interquartile range (IQR: 25th–75th percentiles) in the following.

#### Detection and exclusion of licking periods

In 18 of the 99 recording sessions in the dataset KI, mice could lick to obtain water rewards given after the tone stimuli. Licking requires motor action which could influence firing patterns. Licking also occurred outside the reward window and could thus not only affect response epochs but also spontaneous epochs. To improve comparability of patterns across sessions, licking periods were detected and both spontaneous and response epochs overlapping with licking periods were excluded. If available, licking was detected using a piezoelectric sensor. Otherwise licking was detected as artifacts in LFPs (LFPs down-sampled to 500 Hz with a polyphase filter) using independent component analysis. Lick periods invariably presented as a single independent component displaying a characteristic, rhythmic, saw-tooth like pattern with close to uniform weight contribution across the electrodes of a probe, i.e. the lick artifacts were similarly strong across electrodes. Periods of lick artifacts in the lick component were detected using a semi-automatic procedure which involved manually setting an amplitude threshold for each recording, detecting threshold crossings, and manual curation that allowed deletion and inclusion of false positive and false negative detections, respectively.

#### Detection of sleep-like periods

Sleep-like periods^26^ were detected per recording session as illustrated in Supplementary Data Fig. 1c. A mean firing rate vector across all cortical spikes (bin-width: 10 ms) was computed and smoothed with a Savitzky-Golay filter (window width: 11 pts, order: 3) to obtain the mean smooth firing rate vector FR(mean). Two thresholds were defined, ϴgmean, the geometric mean of FR(mean), and ϴtrough = 0.2* ϴgmean. Periods when FR(mean) was below ϴtrough for longer than 5 ms were detected as ‘troughs’. To obtain ‘off periods’, troughs were extended forwards and backwards in time until FR(mean) reached above ϴgmean. Off periods formed the basis of sleep-like activity. Successive off periods closer than 1.5 s were merged. Around merged off periods and individually occurring off periods 1 s and 0.3 s, respectively, were included into the sleep-like periods. All sleep-like periods were reviewed visually and consistently coincided with a flat EMG, indicating absence of movement, and large amplitude, irregular, slow-wave activity in the LFP typical of non-REM sleep and drowsiness^26^. Detection of sleep-like periods was only performed for the dataset KI as sessions of this dataset consistently featured a sufficient number of cortical units to detect collective off periods. Sleep like episodes occupied 6% (IQR: 0–18%) of the time in recording sessions.

#### Epoch selection

##### KI, spontaneous epochs

In the dataset KI, spontaneous epochs were selected as time windows from 3 s before tone onset until tone onset. All epochs containing licking (see ‘Detection and exclusion of licking periods’) and where saturation occurred in the spike band were excluded. Sleep-like epochs were defined as all episodes overlapping with sleep-like periods (see ‘Detection of sleep-like periods’). Sleep-like epochs are only featured in Supplementary Data Fig. 3c–e. All other analyses of spontaneous activity are based on ‘spontaneous active’ epochs occurring with at least 1 s temporal distance to sleep-like episodes. Recording sessions contained 227 (IQR: 172–284) spontaneous active epochs and 19 (IQR: 1–84) sleep-like epochs.

##### IBL, spontaneous epochs

In all the 303 recordings of the dataset IBL (IBL Neuropixels Brainwide Map ^42^ on AWS was accessed on February 2024 from https://registry.opendata.aws/ibl-brain-wide-map.) a 5 min block of spontaneous activity was recorded towards the end of the recordings, after 68 min (IQR: 58–80 min). Here, spontaneous epochs were obtained by splitting the 5 min block into 99 epochs lasting 3 s each.

##### Robustness of results for shorter epoch durations

We selected a pre-stimulus epoch of 3 s, as the inter-tone intervals varied randomly between 5 and 10 seconds, and tone-evoked responses were observed to dissipate within 2 s after tone onset. Analyses based on shorter pre-stimulus epochs of 2 s or 1 s were also performed, but fewer units could be included due to insufficient data (number of spikes) for reliably estimating activity patterns in these shorter epochs. However, when using shorter epochs, results remained qualitatively consistent, validating the robustness of our findings.

##### KI, response epochs

In the dataset KI, epochs used to compute response traces were selected as time windows starting 2 s before tone onset until 0.7 s after tone onset. All epochs containing licking, sleep-like periods, and where saturation occurred in the spike band were excluded. Recording sessions contained 188 (IQR: 86–228) response epochs. The number of response epochs per recording was lower than the number of spontaneous active epochs per recording because licking disproportionately occurred during response time windows, which coincided with reward presentation in some recordings (see ‘Detection and exclusion of licking periods’).

#### Waveform classification

A maximum of 2,000 randomly selected 2.8 ms waveforms per unit were extracted from the spike band. 82 samples points (–1.4 ms and +1.4 ms) around each spike time provided by Kilosort2 were collected for each waveform. The mean waveforms per unit were obtained by averaging across all collected waveforms per unit. Each mean waveform was converted from 16-bits analog values (i) to voltage values (V) as follows:

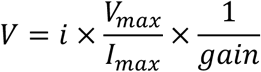

With Vmax = 0.6 V, Imax = 512 bits, and gain = 500, where the factor (Vmax) ÷ (Imax × gain) is the least significant bit. The mean waveforms were then interpolated by a factor of 1000 (1D linear interpolation) and baseline corrected. A custom script was used to detect and compute the main peaks, amplitude values, the polarity of the waveform, the main slopes and amplitude ratio on the interpolated mean waveform per unit. Only the peak to trough duration was used for waveform classification. The valley observed in the peak to trough duration distribution (Fig. 1h) was used to divide units into nw units (peak to trough duration < 0.38 ms), ww units (peak to trough duration > 0.43 ms), and unclassified units (0.38 ms ≤ peak to trough duration ≤ 0.43 ms) according to previous studies^29^.

#### Extraction of spontaneous activity metrics

Spontaneous activity metrics were extracted from spontaneous epochs (see ‘Epoch selection’). For each unit, the three metrics characterizing firing rate, burstiness, and memory were first calculated per epoch. The firing rate metric log10FR was computed as the decadic logarithm of the number of spikes in an epoch divided by epoch duration (3 s). The burstiness and memory metrics were based on inter-spike intervals (ISIs) and defined according to^27^. In brief, burstiness ‘B’ can be understood as the coefficient of variation of ISIs normalized to a range between –1 (completely regular; std(ISI)<<mean(ISI)) and 1(maximally bursty; std(ISI)>>mean(ISI)) and was calculated as:

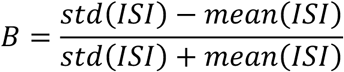

with mean(ISI) and std(ISI) being the mean and standard deviation of ISIs, respectively. Memory ‘M’ was defined as the Pearson correlation coefficient (PCC) between successive ISIs:

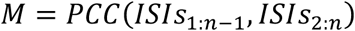

Burstiness and memory were only computed in epochs with at least six spikes available (corresponding to at least four ISIs). Units with fewer than 12 epochs with at least six spikes were excluded from further analyses. To characterize a unit’s firing pattern, each of the three metrics were averaged across epochs. Note that calculating burstiness and memory per epoch before averaging has the advantage that the distribution-based burstiness and the sequence-based memory can both be faithfully computed. The metric ‘local variation’ (LvR) proposed by Shinomoto et al.^43^ was designed to mitigate vulnerability of ISI-based irregularity metrics to slow fluctuations in firing rates when computing ISI-based metrics over longer stretches of time. Yet, given the brief and stationary (stimulation free) character of the spontaneous epochs in the dataset KI, burstiness and memory are a more informative variable choice than LvR as these metrics allow to address the distributional and sequential character of bursting independently of each other.

#### Extraction and post-processing of responses

Responses were extracted from response epochs, which spanned an interval from 2 s before the stimulus (‘prestim-window’) to 0.7 s after the stimulus (‘poststim-window’; see ‘Epoch selection’).

##### Extraction of peri-stimulus time histograms (PSTHs)

For each unit, spikes times relative to tone onset were collected from all its response epochs to construct a firing rate PSTH (bin width: 0.5 ms; Fig. 5a). The PSTH was smoothed by convolving it with a gaussian kernel (standard deviation of the kernel: 10 ms, width of convolution window: 80 ms). Units with an overall average firing rate below 0.1 Hz across all response epochs or an average firing rate below 0.1 Hz across all prestim-windows were excluded from further analyses. To obtain a trace reflecting relative stimulus-induced rate changes, each unit’s smoothed PSTH was z-scored to its prestim-window through subtracting the mean and dividing by the standard deviation across the prestim-window. Response traces were defined as the smooth z-scored PSTHs extending from stimulus onset to 0.65 s post onset.

##### Dimensionality reduction of response traces

To facilitate later classification of responses (see ‘Self-organizing maps’ and ‘Hierarchical clustering analyses’ below) the dimensionality of the response traces was reduced using principal component analysis (PCA). Response traces from all units (both nw and ww units pooled) were collected into a data matrix (dimensions: [number of units, number of response trace time-points]). Each time-point in the data matrix was mean-centered. PCA was performed on the time-points. The top 8 principal components, explaining together over 95% of the variance, were retained. The dot product of a unit’s original response trace and the principal components resulted in 8 scores summarizing the response trace of a unit.

#### Self-organizing map analysis

A self-organizing map (SOM) is an unsupervised machine learning algorithm^44^ used here to summarize a set of n-dimensional feature vectors into a two-dimensional grid of nodes. Each node represents an n-dimensional prototype vector (visualized as a hexagon). After training the SOM, similar prototype vectors become neighbors on the SOM grid. Separate SOMs were constructed for the following data selections of the dataset KI: (i) spontaneous activity of ww units, (ii) spontaneous activity of nw units, (iii) responses of ww units, and (iv) responses of nw units.

##### Input features

To obtain input for training the SOM, each unit *j* was represented by a feature vector. For spontaneous activity (i and ii), the feature vector *xj* consisted of the three spontaneous firing metrics: *xj = [log10FR, burstiness, memory]*; see ‘Extraction of spontaneous activity metrics’ above. For response activity (iii and iv), the feature vector comprised eight principal component scores (PCSs) summarizing the response trace of each unit: *xj=[PSCj,1,PCSj,2,…PCSj,8]*; see ‘Extraction and post-processing of responses’ above. Feature vectors were collected from all units in the respective data selection. Each feature was standardized by subtracting the mean and dividing by the standard deviation across all included units.

##### SOM architecture and initialization

The number of nodes in the SOM was manually determined to balance a detailed representation of the input space (requiring more nodes) with the intent to preserve neighborhood relationships and maintain a compact visualization (requiring fewer nodes). The shape (x-y dimensions) of the SOM and initialization of nodes were determined using the linear initialization method suggested by Kohonen (2001)^45^. This involved performing a PCA on the input feature vectors and setting the height and width of the SOM proportional to the ratio of the two largest eigenvalues.

##### SOM training

The SOM was trained using the batch training method and the neighborhood functions provided by Kind and Brunner (2013)^46^. In brief, each feature vector (summarizing a unit’s characteristics) was first assigned to the SOM node with the closest prototype vector in Euclidean space, referred to as the best matching node (BMN). The prototype vectors of the BMUs and their neighboring nodes were then updated to more closely resemble their assigned input feature vectors, using a Gaussian neighborhood function. Over the 200 training iterations, the neighborhood radius was gradually reduced, leading to more localized and subtle updates to the prototype vectors. After the SOM was trained, each unit was assigned to (i.e. represented by) its BMN.

##### Projection of the dataset IBL on SOMs of the dataset KI

IBL data was projected on the SOM trained on the respective subset of dataset KI by extracting the same features from units of the dataset IBL and standardizing them to the mean and standard deviation of the features of the respective subset of dataset KI. As for the dataset KI, a BMN on the SOM derived from the dataset KI was then assigned to each IBL unit.

##### Advantages of SOMs

The SOM here serves as the first step in a two-step clustering procedure (with the second step detailed in the section ‘Hierarchical clustering analyses’, below). Each prototype vector can be viewed as representing a cluster of similar units. This coarse graining into prototype vectors makes subsequent clustering less sensitive to outliers. Additionally, SOMs are well-suited for heuristic approaches like ours, as they offer a compact visualization of similarity structures and can accommodate and elucidate nonlinear relationships between features (metrics). Furthermore, new data can be readily projected on existing SOMs (as done here for the dataset IBL), which increases reproducibility and comparability of results across datasets.

#### Hierarchical clustering analyses

Hierarchical trees were computed for three types of data: (i) SOM nodes (e.g., Fig. 2c, detailed illustration in Supplementary Data Fig. 3b), (ii) PFC flatmap ROIs (e.g., Fig. 4c), and (iii) enrichment matrices (e.g., Fig. 2g), using Ward’s agglomerative hierarchical clustering algorithm^47^.

##### Feature selection and standardization

Prior to clustering, all features were standardized by subtracting the mean and dividing by the standard deviation across samples. The features used for constructing the hierarchical tree varied by type of data: SOM nodes were represented by prototype vectors (see ‘Self-organizing map analysis’), and PFC flatmap ROIs and enrichment matrices by enrichment profiles (see ‘Enrichment statistics’). In the case of enrichment matrices, the hierarchical tree was used solely to order the matrix’ rows for better visualization.

##### Clustering

For SOM nodes and PFC flatmap ROIs, the hierarchical tree was cut at a specific level to define clusters representing categories of spontaneous/response activity (e.g., Fig. 2c and Fig. 5c) or modules of spontaneous/response activity (e.g., Fig. 4c and Fig. 5f), respectively.

##### Categorization of unit activity

Specifically, in the case of spontaneous/response categories obtained by clustering the SOM nodes, units inherited their category from their BMN (best matching node). Repeating analyses using various numbers of unit activity categories, i.e. SOM clusters (ww: 5–10 categories; nw: 4–8 categories), led to qualitatively similar results, validating the robustness of our approach.

##### Determining a suitable number of clusters

The number of clusters (i.e. implicitly the threshold for cutting the hierarchical tree) was manually determined according to the Thorndike criterion^48^ and the Dunn index^49^, aiming for a balance between cluster compactness and separation (as e.g. shown in Supplementary Data Fig. 3a) and for summarizing the data with the smallest number of clusters that still aligned with both criteria.

#### Stability analyses

To assess the validity of unit categories, we tested how stable unit categories were when subdividing data into blocks and evaluating unit category per block.

##### Obtaining spontaneous unit categories per block

Obtaining the categories of a unit per block involved three steps: (i) allocating epochs to each block, (ii) calculating spontaneous metrics per block, and (iii) categorizing the unit in each block. (i) In experimental settings where there was an inherent block structure, spontaneous epochs belonging to a certain inherent block were used. Experimentally defined blocks contained 50 or 100 epochs. Blocks of 100 epochs were split in two. In experimental settings without block structure, epochs were allocated to blocks of around 50 epochs. There were 5 (IQR: 3–5) blocks available per recording. Epochs with licking or sleep-like activity (see ‘Epoch selection’) were discarded from each block. For each unit, blocks with fewer than 12 epochs with at least six spikes were excluded from further analyses. Per unit, blocks analyzed contained 36 (IQR: 23–49) epochs for ww units and 43 (IQR: 29–51) epochs for nw units, respectively. (ii) For each unit, metrics were extracted per epoch and then averaged per block, as detailed in ‘Extraction of spontaneous activity metrics’. (iii) For each unit, the three spontaneous metrics (features) describing a block were standardized to the mean and standard deviation of the respective original dataset used to compute the reference SOM (see ‘Self-organizing map analysis’). Block-wise ww unit data was standardized to and projected on the SOM derived from the original (full epoch) ww dataset. Block-wise nw unit data was standardized to and projected on the SOM derived from the original (full epoch) nw dataset. After standardizing, the BMN of each unit was identified on the respective SOM. The block-specific category of a unit was inherited from the BMN (see ‘Hierarchical clustering analyses’).

##### Assessment of overall stability per transition

For each unit *u* and transition *t* from one block to the next, we calculated a transition matrix *Mu,t* (dimensions: *k × k*; where *k* is the number of categories available for a dataset, i.e. *k=8* for ww units, and *k=5* for the nw units). This binary matrix contains exactly one non-zero entry at the row corresponding to the category the unit had in the current block and the column corresponding to the category the unit transitioned to in the next block. For each transition, the unit-wise transition matrices *Mu,t* were summed across all units, resulting in a *k × k* transition matrix *Mt* for each transition *t* (*t=1* corresponds to the transition from block 1 to block 2, *t=2* corresponds to the transition from block 2 to block 3, etc.). Each entry *mi,j,t* represents the number of times units of a certain category *i* in a block transitioned to a certain category *j* in the following block during a transition *t*. We defined the stability value per transition as the sum of the diagonal elements in *Mt* divided by the sum of all the entries in *Mt*. The stability value thus expresses the fraction of units that retained their category in a transition. In Fig. 2e the stability value is contrasted with the chance stability expected from the marginal distributions of *Mt*.

##### Assessment of stability per category

To assess how stable individual categories were and which categories preferentially transitioned to which other categories, we summed transition matrices *Mt* across transitions to obtain an overall *k × k* transition count matrix *M* (Supplementary Data Fig. 3d, first panel) and analyzed various matrices derived from *M*. The transition probability matrix *P* was obtained by dividing the entries of a row *i* in *M* by the sum of the row entries, such that each row sums to 1 (Supplementary Data Fig. 3d, third panel, rows and columns reversed for visualization). Here, each entry *pi,j* gives the probability of transitioning to category *j* in the next block given category *i* was assigned in the current block. Evaluating stability from *P* has the disadvantage that categories more abundantly present in the dataset will appear more stable purely by chance. We therefore also considered the observed-to-expected ratio matrix *O* (Supplementary Data Fig. 3d, last panel). For this we first defined an expected transition count matrix *E* derived from the marginal distributions of *M* (Supplementary Data Fig. 3d, second panel). An entry *ei,j* in *E* is defined as *(Ri×Cj)/n*, with *Ri* being the row total count of category *i*, *Cj* the column total of category *j,* and *n* the grand total of counts of *M. O* was obtained by dividing *M* by *E*. Each entry *oi,j* gives thus the ratio of the observed count and the expected count. This display accentuates the stability of overall small categories yet underemphasizes the stability of large categories.

##### Coincidence coefficient

To overcome the interpretative limitations of *P* and *O*, we developed the coincidence coefficient matrix *X* (Fig. 2f). We defined the coincidence coefficient *xi,j* as:

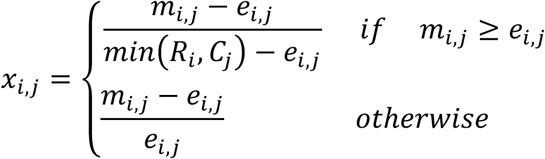

The coincidence coefficient ranges from –1 to 1. A value of –1 indicates that a transition never happens, 0 indicates that the transition happens as often as expected by chance, and 1 indicates that the transition happens as often as possible (relative to the size of the smallest category considered for the transition, i.e. *min(Ri,Cj)*).

#### Enrichment statistics

The need to control for sampling differences between groups (e.g., groups of animals undergoing different behavioral settings) necessitated a statistical approach beyond fractional composition to determine whether a certain attribute *c* (e.g., a unit activity category) is disproportionately frequent or rare in a statistical population *r* (e.g., all units in a certain brain region). To avoid biases introduced by group-specific sampling, we developed an enrichment statistic that uses group-level shuffling (see Supplementary Data Fig. 3c for an illustration of the procedure described here).

##### Obtaining the coincidence matrix X

For each group *g*, it is counted how often an attribute *c* appears within a population *r*, generating a group-specific coincidence count matrix *Xg* (dimensions: *n × m*; where *n* is the number of populations and *m* is the number of attributes). Each entry *xr,c,g* represents the number of times *c* occurred in *r* for group *g*. Summing these matrices across all *k* groups produces the overall *n × m* coincidence matrix *X*, with *xr,c = xr,c,1+ xr,c,2+….+ xr,c,k*.

##### Obtaining the surrogate matrix S

To estimate the coincidences expected by chance while controlling for group sampling biases, *r*-*c* combinations are shuffled (dissociated and randomly re-associated) within each group, generating a surrogate dataset *j*. The shuffling is repeated 1,000 times, creating 1,000 surrogate datasets. For each surrogate dataset, the process of obtaining *X* (described above) is reused, leading to a group-specific surrogate coincidence matrix *Sg,j* and through summing over groups to an overall matrix *Sj*. Ultimately, all 1,000 *Sj* are combined into a *n × m × 1,000* surrogate matrix *S*.

##### Calculating the E-score

The E-score *er,c* for each *r*-*c* combination is calculated by standardizing the original coincidence count *xr,c* to the distribution of the 1,000 surrogate counts in *sr,c,1:1000*:

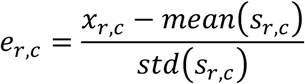

E-scores for all *r*-*c* combinations are collected in the *n × m* enrichment matrix *E*.

##### Significance of enrichment

The significance of an enrichment (p-value, *pr,c*) is obtained from the fraction of surrogate counts *sr,c,1:1000* falling below (*fr,c,-*) or exceeding (*fr,c,+*) the original coincidence count *xr,c*: *pr,c = 1*– *max(fr,c,-,fr,c,+)*.

##### Group-bias control employed for specific types of data

Category enrichment statistics based on (sub)regions and PFC flatmap ROIs (e.g., Fig. 2g, Fig. 4c, and Supplementary Data Fig. 3g) were group-controlled for behavioral settings. The layer-based category enrichment statistic (Supplementary Data Fig. 3f) and the response category vs spontaneous category enrichment statistics (Fig. 5j) were controlled for combinations of behavioral settings and brain (sub)regions, i.e., each combination of a sub(region) and a behavioral setting constituted a separate group *g*.

#### PFC flatmap projection and parcellation

For a compact 2D representation of the PFC, we used the flatmap kindly provided by Gao et al., (2022)^10^. Each unit was positioned on the flatmap by projecting the 3D stereotaxic coordinates (AP, ML, DL) from the unit’s recording site (where the unit’s waveform exhibited the highest amplitude) into the corresponding (u,v) coordinates of the flatmap.

##### Parcellation into ROIs of similar unit count

To generate statistically comparable regions of interest (ROIs) that respected subregion boundaries, we parcellated cytoarchitecturally defined subregions of the PFC flatmap into dataROIs, each containing a similar number of units (Supplementary Data Fig. 7a). This parcellation was based on deep layer (L5–6) units from the dataset KI, aiming for approximately 200 units per dataROI. The subregion FRP, which contained fewer than 200 units in total, was not subdivided further. All other subregions were subdivided as follows: The number of dataROIs *n* for each subregion was determined by *n=⌊(unit count in a subregion)/200⌋*. The (u,v) coordinates of the units contained in the subregion were then clustered into n groups using a constrained K-means algorithm^50^ . with the target range of 160–240 units per dataROI serving as the size constraint for each cluster. The subregion was then parcellated using a Voronoi diagram^51^ based on the cluster centers, such that a ROI was defined as the space closet to a cluster center.

#### ZETA-test

To test for single unit responsiveness to auditory stimulation and estimate the onset of the responses we used the parameter-free Zenith of Event-based Time-locked Anomalies (ZETA) test^39^. We analyzed the 700 ms response time windows following tone onset and obtained p-values and response onset time (the time after which half of the response peak is reached) for each unit. Units with a p-value < 0.01 were considered responsive to the auditory stimulus.

#### Linear mixed-effect regression models

To estimate the effect of neuron type on each spontaneous firing metric (log10FR, burstiness and memory), we fitted linear mixed-effect (LME) regression models with unit type (categorical variable: ww or nw) as a fixed effect and mouse as random effect^52^:

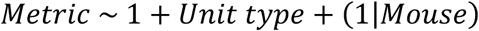

Significance of the effects and interactions was assessed with post hoc F-tests^53^ using the Kenward-Roger method for degree of freedom adjustment^54^ and Tukey’s method for multiple comparison correction. We repeated this LME analysis to estimate the effect of epoch type (categorical variable ‘active’ or ‘sleep-like’) on each firing metric with the following model:

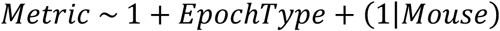

#### Correlation between firing metrics

To estimate the correlation between firing metrics, we fitted LME models for each pair of metrics (Metric1 and Metric2) with mouse as random effect:

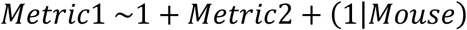

The correlation between any two variables is reported as the marginal coefficient of determination^55^.

## Data and Code availability

The data from the 99 recorded Neuropixels sessions used to generate all main and supplementary figures is available for download in Neurodata Without Borders (NWB) format on DANDI Archive: https://dandiarchive.org/dandiset/000473 and [dandi link will be inserted upon publication].

The code used to preprocess the data and generate all manuscript figures is available at: https://github.com/hejDMC/pfcmap The following open-source software and toolboxes were used: **Python**: matplotlib, netgraph, numpy, scipy, shapely, sklearn, SOMz; **Julia**: LightXML, MixedModels, Statistics, StatsBase, StatsModels, StatsPlots; **R**: emmeans, lme4, MuMIn, pbkrtest.

## Acknowledgments

We thank Le Gao and Jun Yan for sharing the flat map code and anatomical hierarchy score data; Gaelle Chapuis, Olivier Winter, and other IBL members for their technical help and early access to the passive IBL dataset. Alexander Wolthon for his technical help in handling our mouse colony. This work was supported by Wallenberg Scholar program (Knut and Alice Wallenberg Foundation), a KAW project grant (Knut and Alice Wallenberg Foundation), the WennerGren foundation, Hjärnfonden, a NARSAD Young Investigator Grant (Brain & Behavior Research Foundation) and a Stratneuro Postdoctoral grant.

## Author contributions

Conceptualization: P.L.M., K.H., and M.C.

Methodology: P.L.M. and K.H.

Investigation: P.L.M., M.S., F.J., E.M., N.G., R.Y., H.P., F.W.

Data analysis: P.L.M. and K.H.

Project Design and Supervision: M.C.

Visualization: P.L.M., K.H., and M.C.

Writing: P.L.M., K.H., and M.C.

All authors discussed and commented on the manuscript.

## Supplementary Data

**Supplementary Data Fig. 1.**
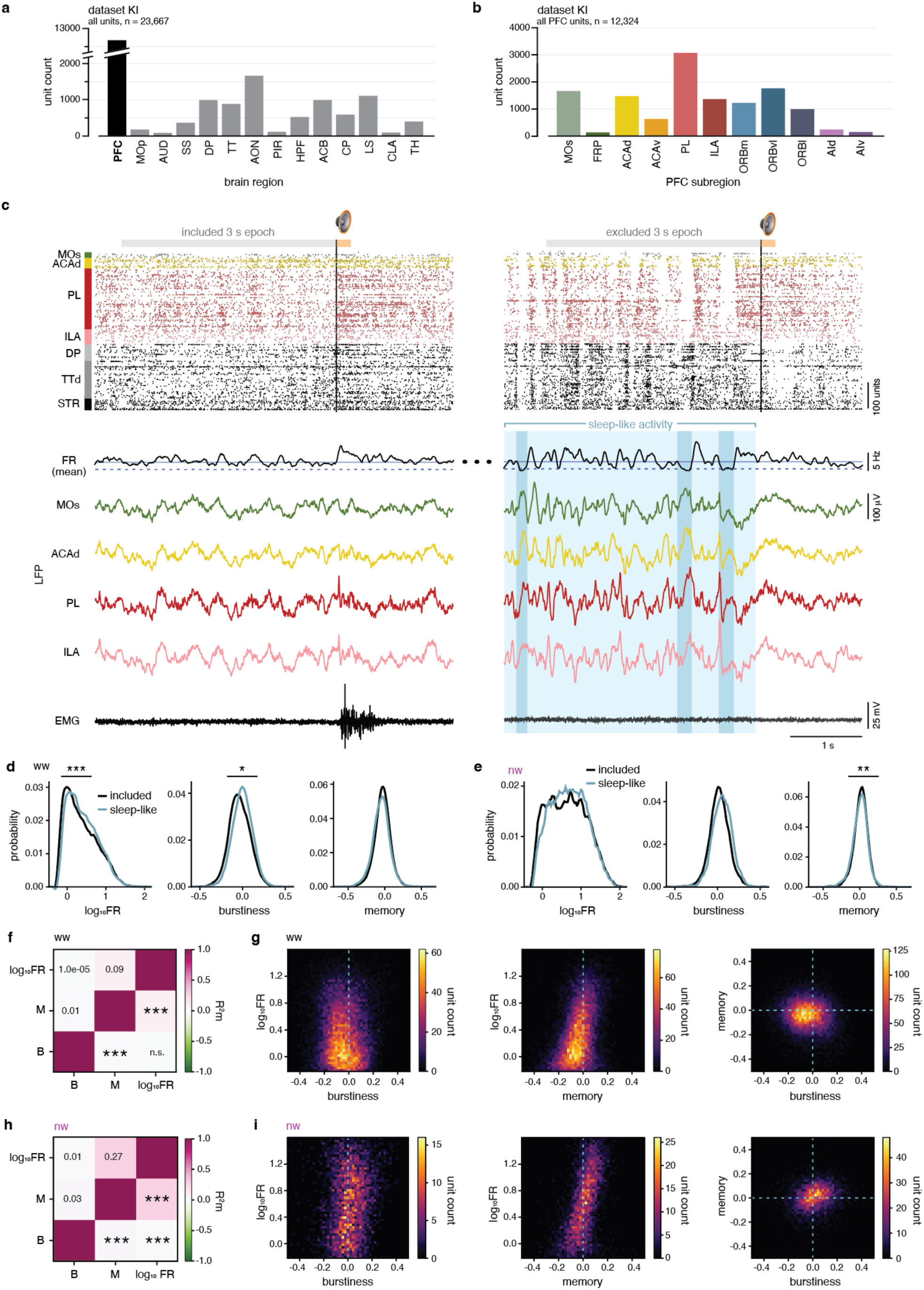
The dataset KI, with description of data selection and correlations of firing metrics. **a,** Count of high-quality, manually sorted, single units included in the current dataset (dataset KI) per brain region. **b,** Count per PFC subregion of all single units included in the dataset KI. **c,** Excluding ‘sleep-like’ periods. Gray horizontal bars indicate 3 s epochs of spontaneous activity directly preceding tone presentation (200 ms, orange bar); black vertical line: tone onset. The left side exemplifies spontaneous epochs included in analysis; the right side exemplifies spontaneous sleep-like epochs excluded from analysis. Any 3 s epoch of spontaneous activity temporally overlapping with a sleep-like period was excluded. Sleep-like periods were identified by the presence of ‘off’ periods (blue vertical bars). Off periods were defined as drops in FR(mean) below the 20% threshold, and extended forward and backward to when FR(mean) reached above its geometric mean. Off periods are accompanied by high-amplitude LFP waves, and sleep-like periods in general coincide with large-amplitude, highly, variable LFP. Time periods neighboring off periods (light blue shadings) were excluded as well (Methods). *Top to bottom*, Raster plot of the single units recorded with the probe shown in Fig. 1b. FR(mean): the smoothed firing rate averaged across cortical units (bin = 10 ms). Blue solid horizontal line: the geometric mean of FR (mean); blue dashed horizontal line: a threshold corresponding to 20% of the geometric mean of FR(mean). Colored traces of local field potentials (LFPs) from the recording site situated centrally in the respective PFC subregion. Concurrent electromyogram (EMG). **d,** Distributions of log10FR, burstiness, and memory for ww units in included (black) and sleep-like excluded (blue) 3 s epochs of spontaneous activity. * *p* < 0.05, *** *p* < 0.001, mixed-effect regression (Methods). **e,** Same as **d** for nw units. ** *p* < 0.01. **f,** Correlation between firing rate, burstiness, and memory across all ww units. Marginal coefficient of determination (R^2^m) values are shown above the diagonal and the respective p-values are shown below the diagonal. n.s.: non-significant, *** *p* < 0.001, mixed-effect regression. **g,** Two-dimensional histogram showing, across all ww units, the relationship between firing rate and burstiness (left), burstiness and memory (middle), and firing rate and memory (right). **h, i,** Same as **f** and **g** for nw units. *** *p* < 0.001, mixed-effect regression. *Data:* dataset KI, all units across brain (sub)regions and layers, n = 23,667 units; **a** dataset KI, all units across subregions and layers of the PFC, n = 12,324 units; **b** dataset KI, ww units, all brain (sub)regions and layers, included 3 s epochs, n = 19,186 units; **d black**, **f, g** dataset KI, ww units, all brain (sub)regions and layers, excluded 3 s epochs, n = 11,599 units; **d blue** dataset KI, nw units, all brain (sub)regions and layers, included 3 s epochs, n = 4,200 units; **e black, h, i** dataset KI, nw units, all brain (sub)regions and layers, excluded 3 s epochs; n = 2,549 units; **e blue**

**Supplementary Data Fig. 2.**
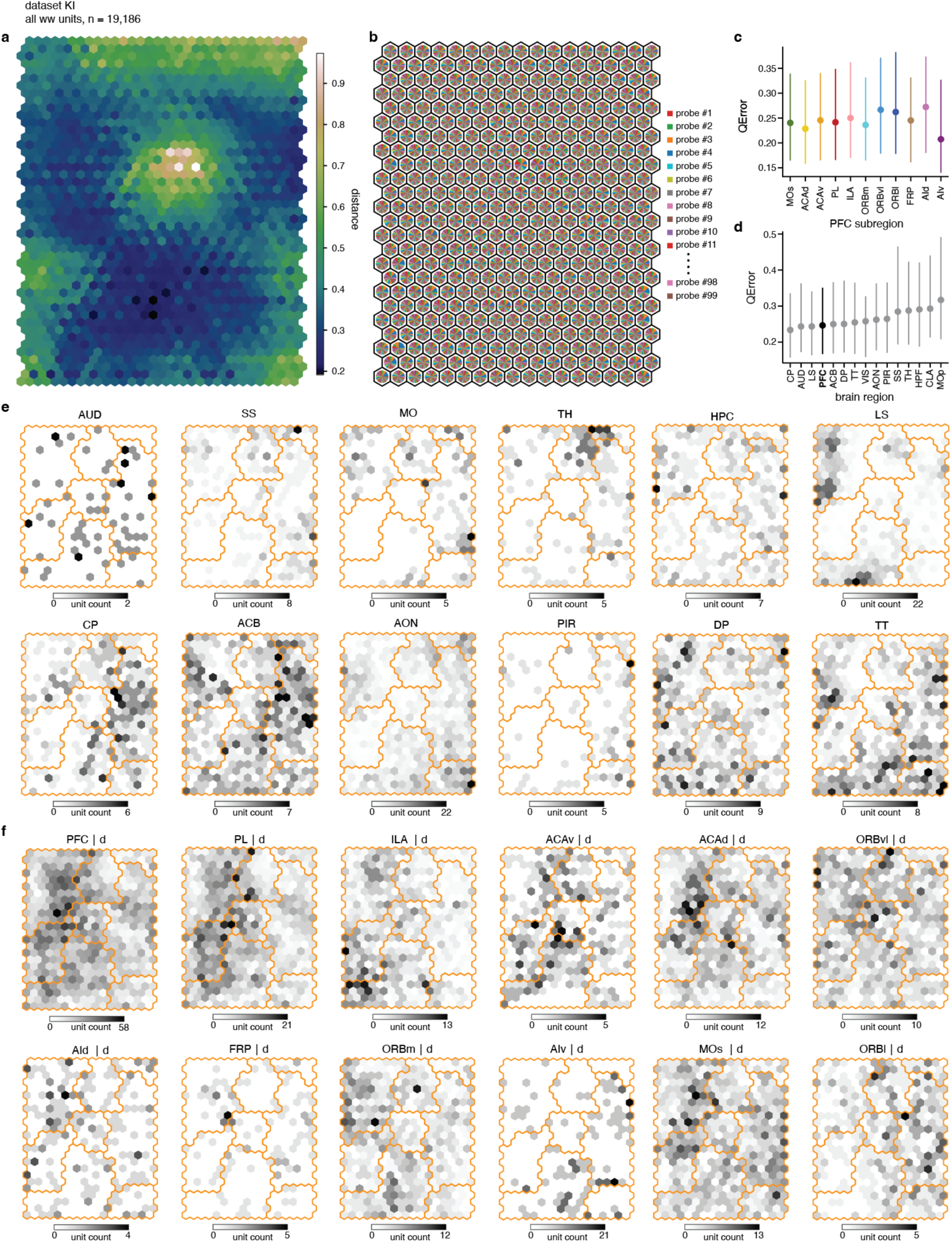
Brain regions occupy distinct territories on the Self Organizing Map. **a,** U-matrix visualizing the distance between neighboring nodes of the SOM trained on ww units (Fig. 2a–e). In the U-matrix, an additional interpolating node is placed between each pair of neighboring (original) nodes. The interpolating node displays in color the Euclidean distance between the pair’s metric vectors. The original SOM nodes are colored based on the average Euclidean distance value of all their surrounding interpolating nodes. **b,** Diversity of units assigned to each SOM node in terms of recording probe. Each pie chart (one pie chart/SOM node) reflects the collection of recording probes sampling units assigned to the node. The pieces of the pie reflect the probes’ respective fraction of ww units. The dataset KI holds 99 recording probes in total. **c,** Quantization error (QError) indicating how well ww units of different PFC subregions were represented on the SOM. The QError is defined as the Euclidean distance between a unit’s metrics vector and the metrics vector of the SOM node the unit is assigned to. A lower QError indicates better representation. Dots: median, vertical lines: 25^th^ to 75^th^ percentile. **d,** Same as **c** but showing QErrors for the PFC overall (black) and the other brain regions sampled (grey). **e,** Numbers of ww units assigned to each SOM node for all sampled brain regions, restricted to deep layers (L5–6) for cortical regions (PFC overall shown in Fig. 2b). Orange contours delineate the unit categories defined in Fig. 2c. **f,** Numbers of ww units assigned to each SOM node for PFC subregions, deep layers (L5–6). *Data:* dataset KI, ww units, all brain (sub)regions and layers, n = 19,186 units; **a, b**, **d, e** dataset KI, ww units, all PFC subregions and layers, n = 10,413 units; **c** dataset KI, PFC ww units, deep layers (L5–6), n = 7,184 units; **f**

**Supplementary Data Fig. 3.**
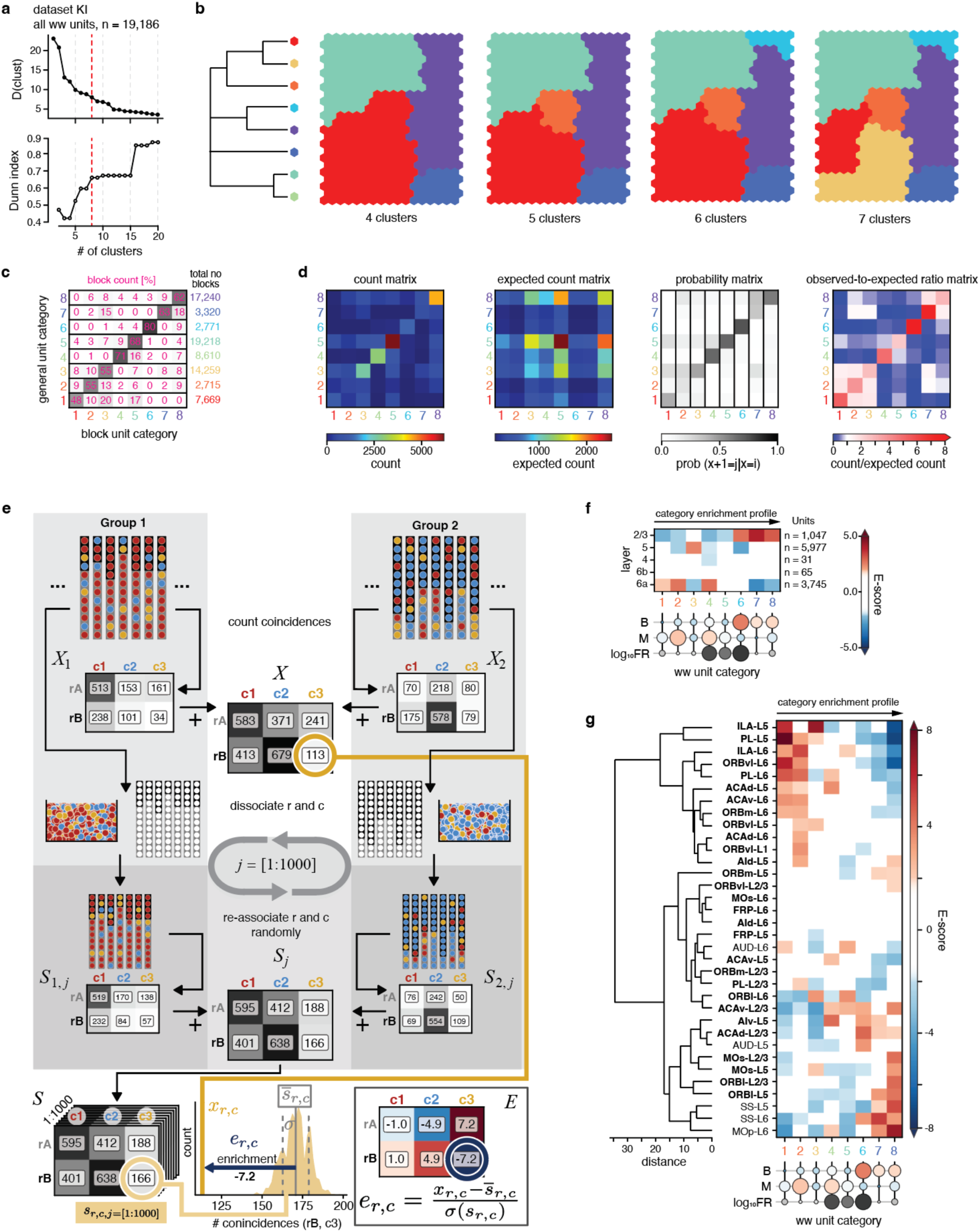
Classification of firing patterns, enrichment statistics, and layer enrichment. **a,** Criteria for determining a suitable number of clusters for partitioning the SOM trained on ww units (Fig. 2c). Red dashed line indicates the eight clusters used. *Top*, Euclidean distance in metric space between the last pair of clusters joined as a function of the number of clusters (categories) defined when hierarchically clustering SOM nodes (read graph from right to left). Lower values indicate more homogenous clusters ^47^. *Bottom*, Dunn index as a function of the number of clusters. The Dunn index is the ratio between minimal between-cluster distance and maximal with-cluster distance, with a higher Dunn index implying more compact and well-separated clusters ^48^. **b,** Hierarchical clustering (ward linkage) of SOM nodes. *Left,* dendrogram displaying the hierarchical relationship of the eight unit categories partitioning the SOM in Fig. 2c (same coloring). *Right,* partitioning of the SOM when opting for four to seven cluster. Coloring follows the respective main branch on the left. **c,** Percentage distribution of the category labels computed per block (obtained when splitting unit data into blocks, Methods) for each unit category (row). Each row sums close to 100%, as percentages were rounded off. For all units belonging to a certain category, all blocks obtained from these units were pooled. Number of blocks per unit category are indicated on the right. **d,** Statistics of category transitions from one block to the next, pooled across blocks. From left to right: count matrix with color indicting how many units that had a certain category in a block (x-axis) transitioned to a certain category in the subsequent block (y-axis); expected count matrix showing the transition counts expected from the marginal distributions of the count matrix (Methods); probability matrix showing the probability of a unit transitioning to a certain category given its category in the current block, with columns summing to 1; observed-to-expected ratio matrix showing the ratio between the observed and expected count for each transition. **e,** Statistical procedure for calculating enrichment scores (E-scores). From top to bottom: Schematic representation of two experimental groups with differential sampling of two brain regions (vertical bars represent recording probes, where grey and black indicate regions rA and rB, respectively) and three unit categories (colored circles). Coincidences of brain regions (*r*) and activity categories (*c*) are counted within each group (*X1* and *X2*) and summed to generate the overall coincidence count matrix across groups (*X*). To estimate the coincidences expected by chance while controlling for group effects, *r* and *c* are dissociated and randomly re-associated for each group, resulting in group-specific surrogate coincidence matrices (*S1,j* and *S2,j*). These surrogate matrices are then summed to obtain an across-group surrogate matrix (*Sj*). The random dissociation, re-association, and coincidence counting process is repeated 1,000 times to generate an overall surrogate matrix, *S*, with dimensions [n regions] x [n activity categories] x 1,000 (bottom left). To calculate the E-score, *er,c*, for a specific activity category *c* in a given region *r* (shown here for region rB and category c3, bottom middle), the original coincidence count *xr,c* (yellow vertical line) is standardized to the distribution of the 1,000 surrogate counts in *sr,c* (yellow shaded area) by subtracting the mean (*sr,c*) and dividing by the standard deviation (*σ*) of *sr,c*. E-scores for all combinations of *r* and *c* are collected in the enrichment matrix *E*. Note that *r* and *c* can represent any attributes (e.g., *r* could also refer to layer or experimental group), and that ‘group’ in this schematic denotes the attribute being controlled for (e.g., ‘group’ could also refer to brain region, when controlling for brain regions, as in **d**). **f,** Category enrichment profiles of different cortical layers. Enrichment profiles were obtained statistically controlling for brain regions (**c** and Methods). Rows are sorted according to hierarchical clustering (ward linkage) of enrichment profiles. Non-significant E-scores are whitened. Circles below summarize the metric composition of each category as detailed in Fig. 2d. **g,** *Right*, Layer-specific category enrichment profiles of cortical (sub)regions. Bold: PFC subregions. Non-significant E-scores are whitened. *Left*, hierarchical tree (ward) derived from the enrichment profiles. *Data*: dataset KI, ww units, all brain (sub)regions and layers, n = 19,186 units; **a**–**d** dataset KI, ww units, all cortical (sub)regions and layers, n = 9,715 units; **d**,**e**

**Supplementary Data Fig. 4.**
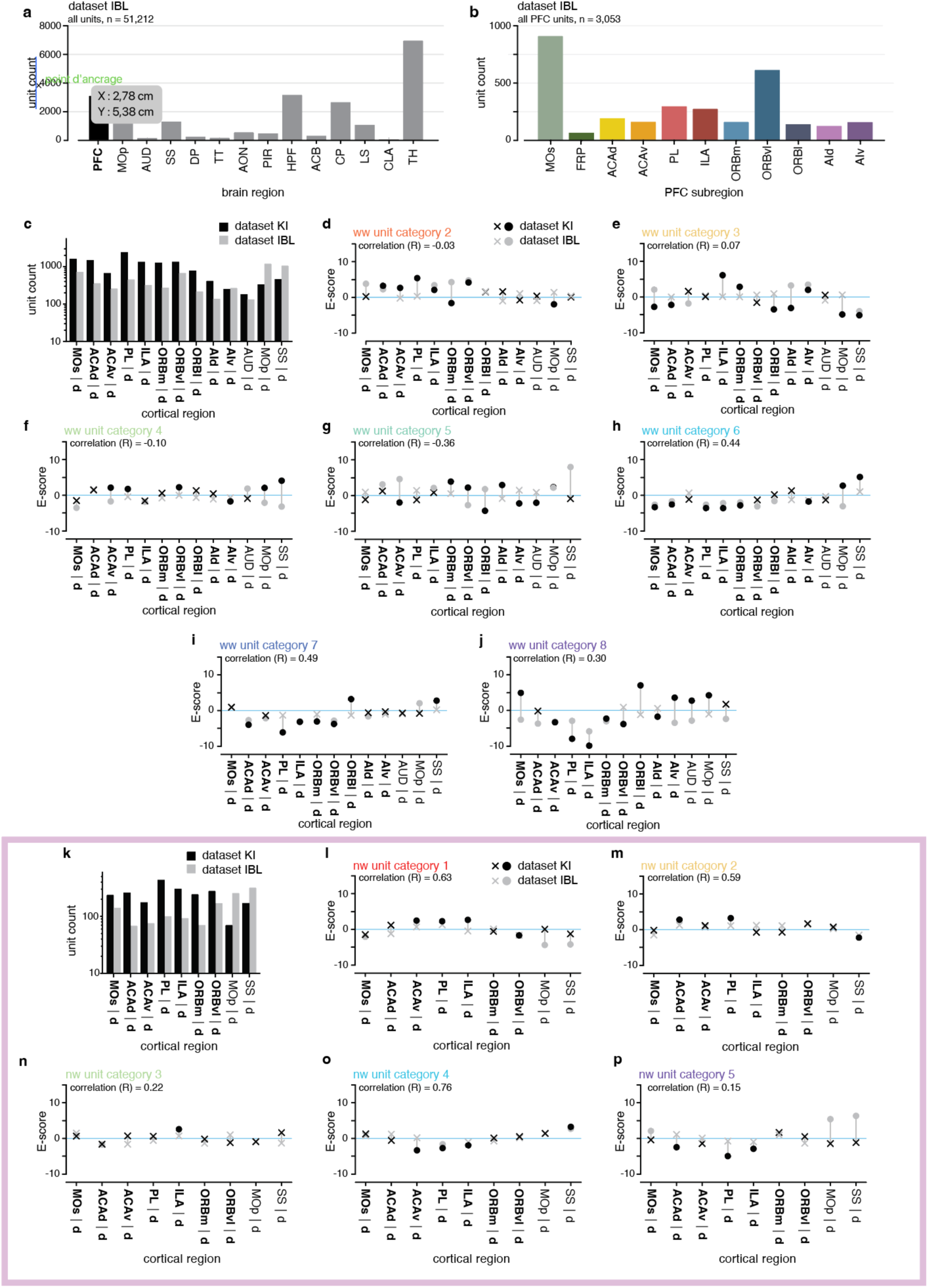
Comparison of category enrichment profiles between dataset KI and dataset IBL. **a,** Count per brain region of all high-quality, manually sorted, single units included in the dataset IBL. **b,** Count per PFC subregion of all single units included in the dataset IBL. **c,** Count per cortical (sub)region of ww units (deep layers, L5–6) in dataset KI (black) and dataset IBL (gray). **d,** Comparison of the enrichment of ww category 2 units in cortical (sub)regions between dataset KI (black) and dataset IBL (gray). Dots and crosses indicate significant and non-significant enrichment, respectively. Correlation between datasets (KI and IBL): R = -0.03. **e to j,** Same as **d**, but for other unit categories. Unit category 3: R = 0.07; category 4: R = -0.10; category 5: R = -0.36; category 6: R = 0.44; category 7: R = 0.49; category 8: R = 0.30. **k,** Count per cortical (sub)region of nw units (deep layers, L5–6) in dataset KI (black) and dataset IBL (gray). **l to p,** Same as **d**, but for nw units (unit categories 1–5). Unit category 1: R = 0.63; category 2: R = 0.59; category 3: R = 0.22; category 4: R = 0.76; category 5: R = 0.15. *Data:*dataset IBL, all units across brain (sub)regions and layers, n = 51,212 units; **a** dataset IBL, all units across subregions and layers of the PFC, n = 3,053 units; **b**

**Supplementary Data Fig. 5.**
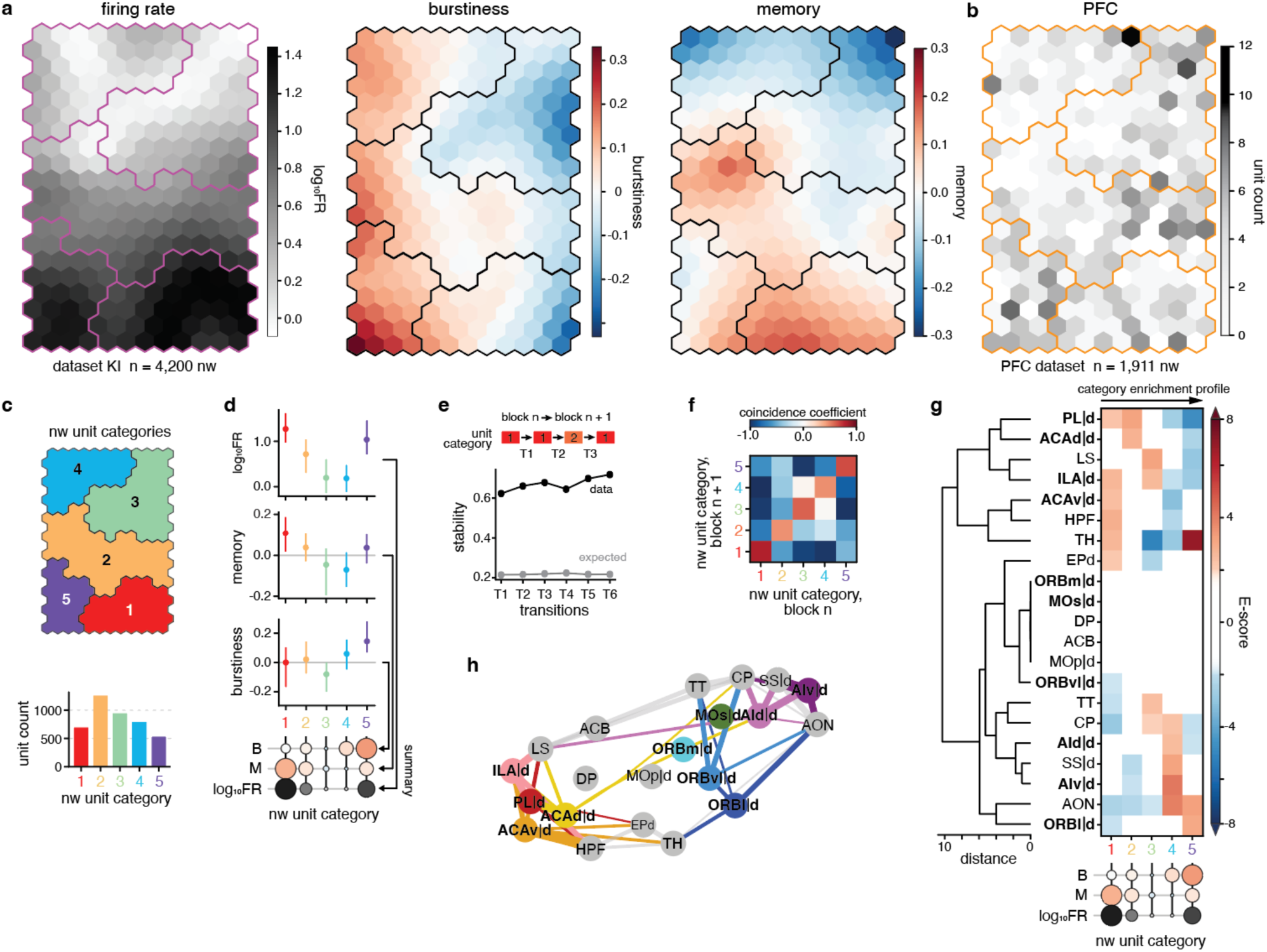
Analysis of category enrichment in brain regions and PFC subregions for nw units. **a,** The component planes of the SOM trained on the three metrics of nw units. Each component plane consists of a hexagonal grid of nodes and displays the respective metric value per node in color. Together, the component planes visualize the combined feature landscape of the SOM. Contours (black/purple) delineate the unit categories defined in **c**. **b,** Number of PFC nw units assigned to each SOM node. Contours (red) delineate the unit categories defined in **c**. **c,** *Top*, Partitioning of the SOM nodes into five unit categories using hierarchical clustering. *Bottom*, number of nw units per unit category. **d,** Summarizing the characteristics of each unit category. Median (dot) and 10th to 90th percentile (vertical line) of metrics across units assigned to each category. Circles below (‘summary’) further summarize the metric composition of each category (filling: median metric value colored according to **a**, radius: relative magnitude of metric value compared to the other categories). **e,** Stability of nw unit categories across time. *Top,* Spontaneous epochs were split into blocks (∼50 epochs/block) and each unit’s category was calculated per block. *Bottom,* Stability (fraction of units retaining their category) from one block to the next (black) compared to stability expected from marginal distributions (grey). **f,** Quantification of the transitions between nw unit categories across all blocks shown as a coincidence coefficient matrix: -1: zero transitions; 0: random; 1: maximal possible number of transitions as derived from the marginal distributions (Methods). **g,** *Right*, Category enrichment profiles (Methods, Supplementary Data Fig. 3c) of brain (sub)regions. Bold: PFC subregions. Non-significant E-scores are whitened (Table 5). *Left*, hierarchical tree (ward) derived from the enrichment profiles. **h,** Graph representation of the data in **g**. Nodes representing brain (sub)regions are arranged according to their first and second UMAP dimension, line width scales with cosine similarity between category enrichment profiles of brain regions (only shown for similarities > 0.1). Bold: PFC subregions. *Data:* dataset KI, nw units, all brain (sub)regions and layers, n = 4,200 units; **a**–**f** dataset KI, nw units, all brain (sub)regions, for cortex restricted to deep layers (L5–6), n = 3,984 units; **g**,**h**

**Supplementary Data Fig. 6.**
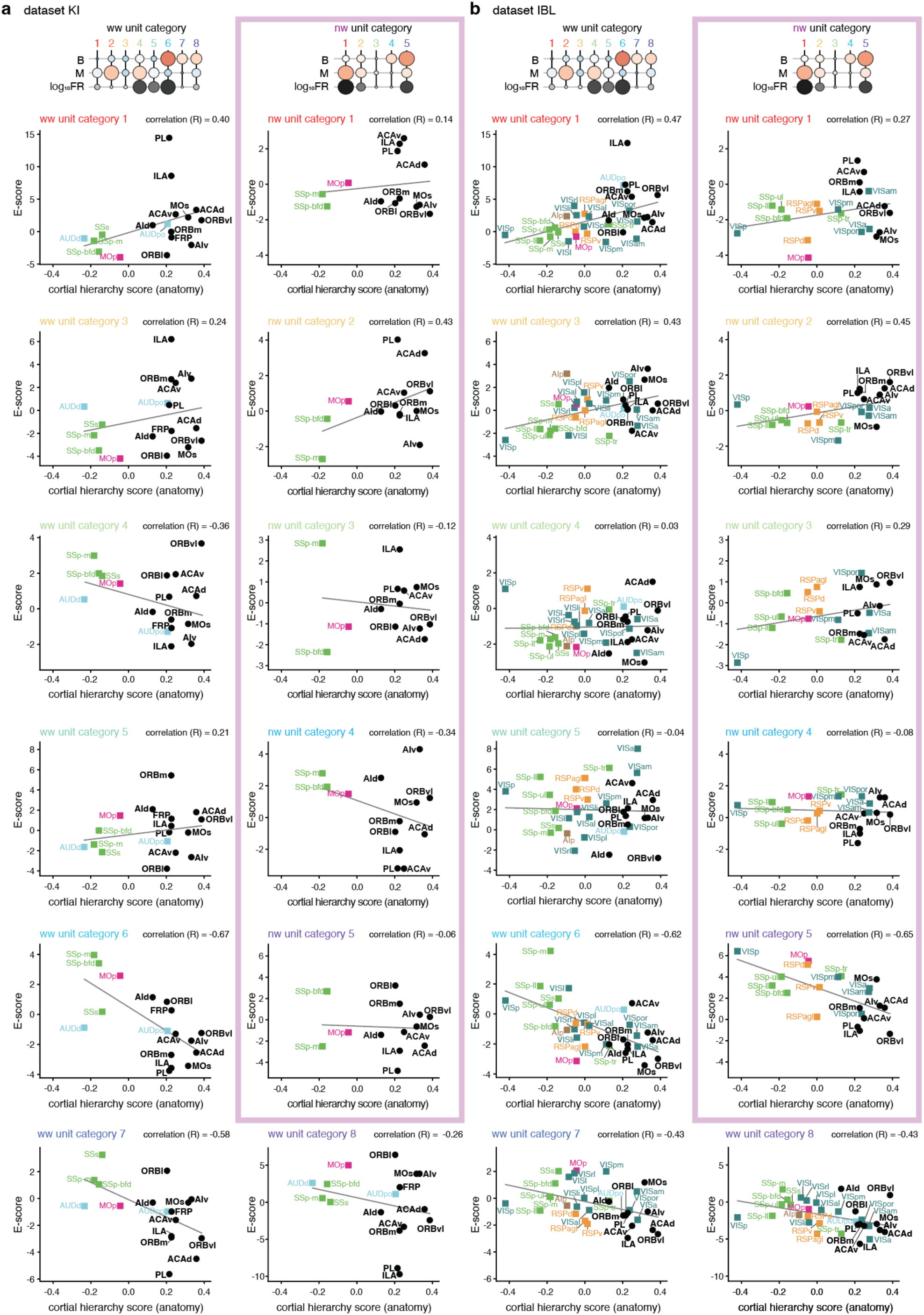
Category enrichment reflects anatomical hierarchy of cortical regions. **a,** Enrichment in unit category vs cortical hierarchy score for dataset KI, ww units: left, nw units: right. Each panel shows the correlation between a unit category and hierarchy score across cortical regions (categories 1, and 3–8 for ww; pink box: 1–5 for nw). Bold: PFC subregions. Pearson correlation: ww unit category 1: R = 0.38; ww category 3: R = 0.25; ww category 4: R = -0.34; ww category 5: R = 0.21; ww category 6: R = -0.68; nw category 1: R = 0.13; nw category 2: R = 0.44; nw category 3: R = -0.14; nw category 4: R = -0.34; nw category 5: R = -0.06. **b,** Same as **a**, but for dataset IBL. Pearson correlation: ww unit category 1: R = 0.47; ww unit category 3: R = 0.43; ww unit category 4: R = 0.02; ww unit category 5: R = -0.03; ww unit category 6: R = -0.62; ww category 7: R = -0.43; ww category 8: R = -0.43; nw category 1: R = 0.26; nw category 2: R = 0.45; nw category 3: R = 0.29; nw category 4: R = -0.08; nw category 5: R = -0.65. *Data:* dataset KI, cortical brain (sub)regions, deep layers (L5–6), n = 10,898 ww units, n = 1,826 nw units; **a** dataset IBL, cortical brain (sub)regions, deep layers (L5–6), n = 7,168 ww units, n = 1,026 nw units; **b** PFC hierarchy scores from Harris et al.^9^.

**Supplementary Data Fig. 7.**
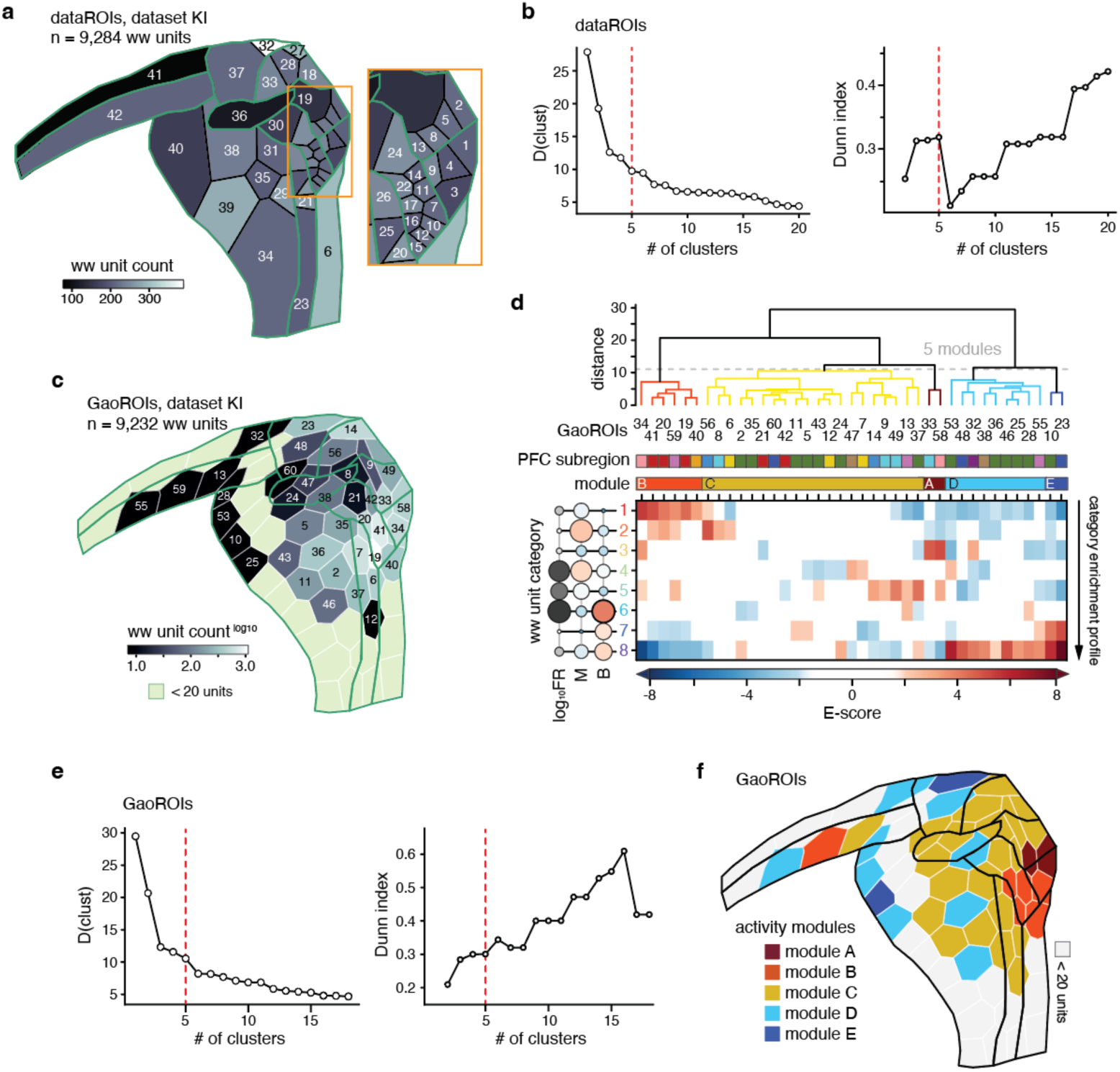
Within PFC sampling, clustering of ROIs, and PFC activity map derived from GaoROIs. **a,** Flatmap of the PFC with the count per dataROI of ww units (deep layers, L5–6) in dataset KI. Green outlines: cytoarchitectural PFC subregions. Each dataROI (black outlines) is identified by an ID number. Box: enlargement of the corresponding frame on the flatmap. **b,** Criteria for determining a suitable number of dataROI clusters to partition the PFC flatmap into modules based on the dataROI’s category enrichment profiles (Fig. 4c,d). Vertical red dashed line marks the five clusters (modules) used. *Left*, Euclidean distance in metric space between the last pair of clusters joined as a function of the number of modules defined when hierarchically clustering the enrichment profiles of dataROI’s nodes (read graph from right to left). Lower values indicate more homogenous clusters ^47^. *Right*, Dunn index as a function of the number of clusters. The Dunn index is the ratio between minimal between-cluster distance and maximal with-cluster distance, with a higher Dunn index implying more compact and well-separated clusters^48^. **c,** Flatmap of the PFC with the count per GaoROI of ww units (deep layers, L5–6) in dataset KI. Each GaoROI (white outlines) is identified by an ID number. Green outlines: cytoarchitectural PFC subregions. GaoROIs with fewer than 20 units (pale green) were not included in analyses. **d,** Clustering of PFC GaoROIs into activity modules based on their category enrichment profiles. *Top to bottom*, hierarchical (ward) tree derived from enrichment profiles; GaoROI ID numbers; PFC subregion identity of the GaoROIs (color-coded as in Fig. 4e); activity modules (A–E); category enrichment profiles of the PFC’s GaoROIs. **e,** Same as **b**, but for GaoROIs. **f,** PFC flatmap with GaoROIs colored according to activity module. Black outlines: cytoarchitectural PFC subregions. *Data*: dataset KI, PFC ww units, deep layers (L5–6), n = 9,319 units; **a** dataset KI, PFC ww units, deep layers (L5–6), n = 9,232 units; **c** to **f**

**Supplementary Data Fig. 8.**
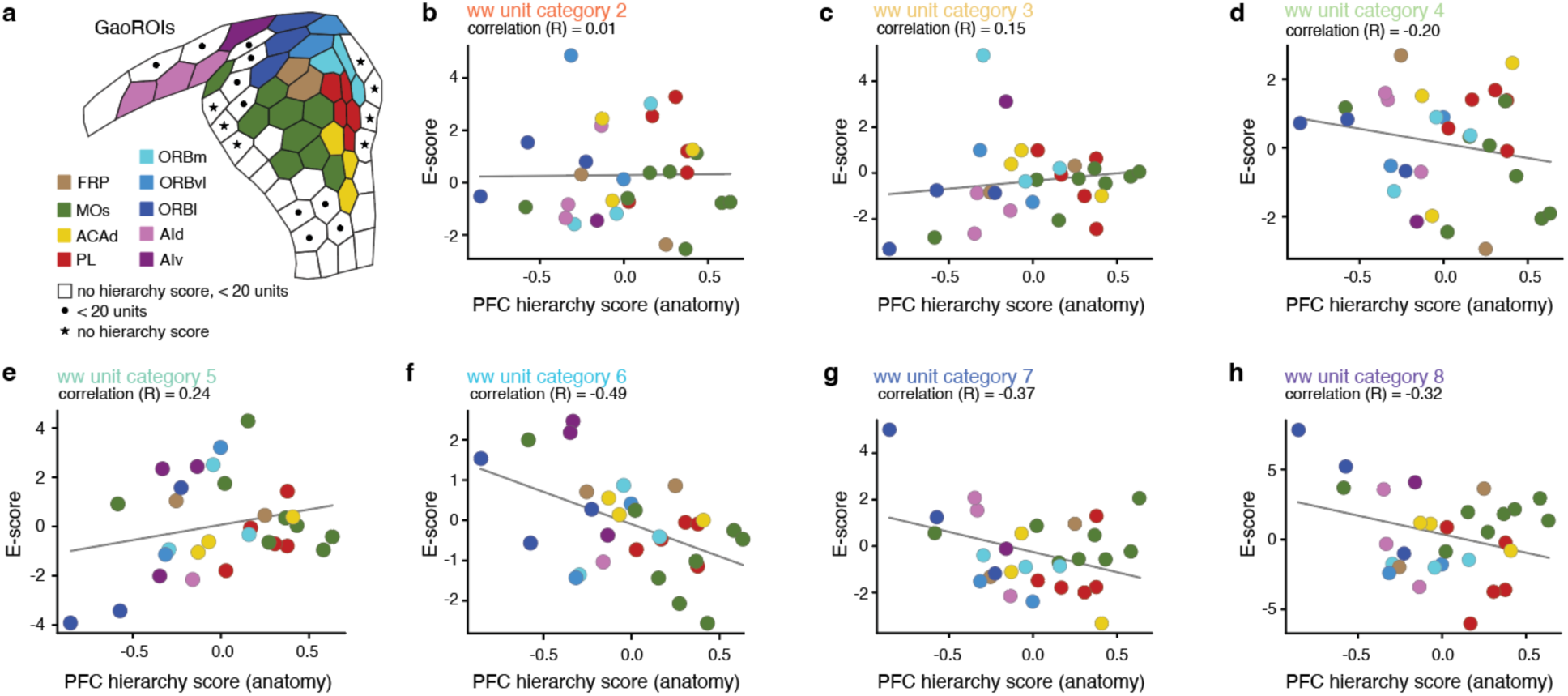
Correlation of enrichments with intra-PFC anatomical hierarchy. **a,** Data availability for the GaoROIs parcellating the PFC flatmap. GaoROIs with fewer than 20 units were not considered for enrichment analyses; GaoROIs with both hierarchy score and category enrichment profile are colored according to the PFC subregion where most of the units were recorded; white: no hierarchy score and too low (n < 20) unit count; black dots: hierarchy score but too low (n < 20) unit count; black stars: sufficient number of units (n ≥ 20) but no hierarchy score. **b to h,** Enrichment in unit category vs PFC hierarchy score for ww categories 2–8. Pearson correlation: category 2: R = 0.02; category 3: R = 0.14; category 4: R = -0.20; category 5: R = 0.24; category 6: R = -0.49; category 7: R = -0.36; category 8: R = -0.31. *Data:* dataset KI, PFC ww units, deep layers (L5–6), n = 9,232 units. PFC hierarchy scores from Gao et al.^10^.

**Supplementary Data Fig. 9.**
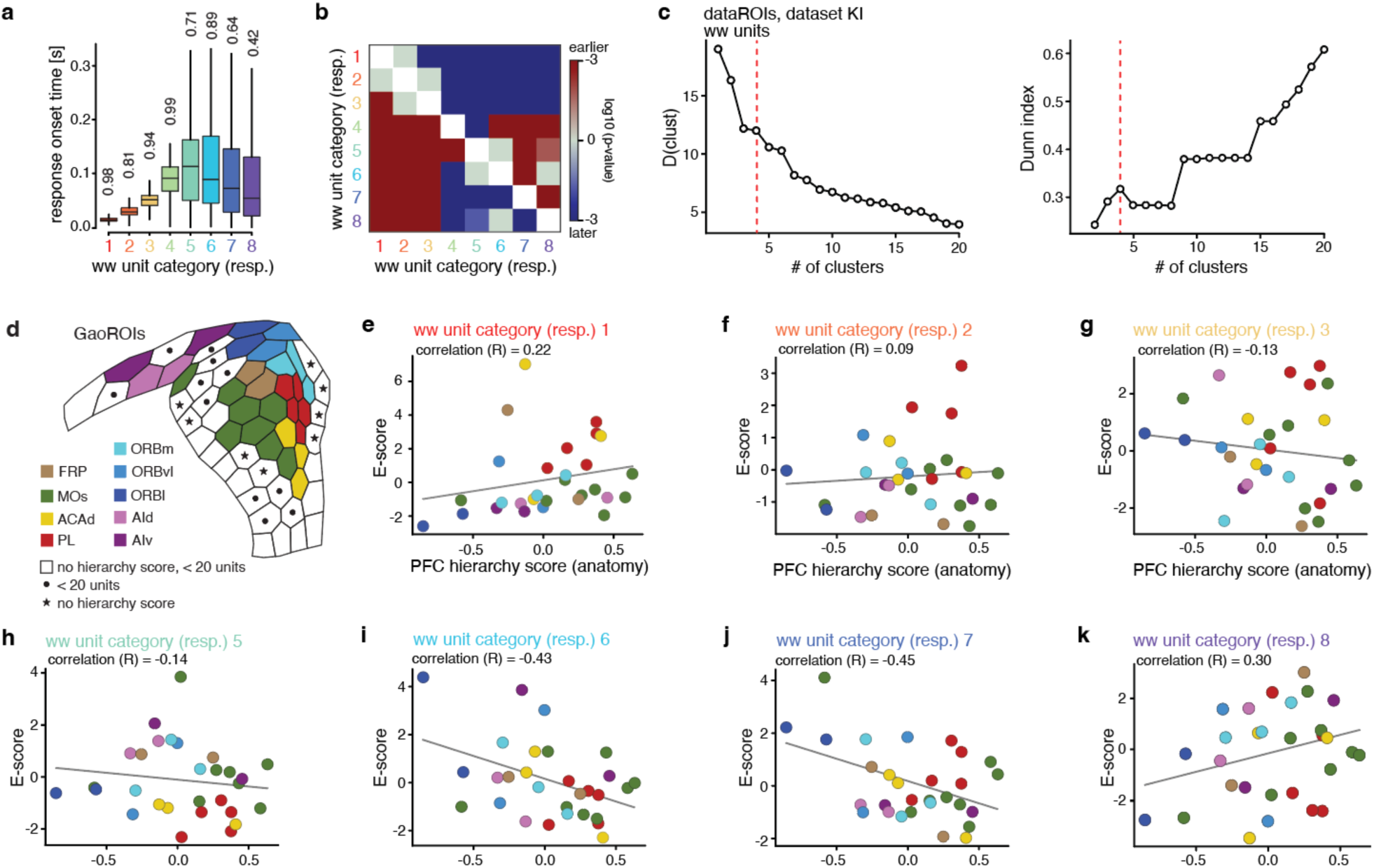
Characterizing ww response categories and correlating their enrichment to intra-PFC hierarchy. **a,** Response onset time (ZETA test^38^) per response category. The whisker plot displays the median (center line), interquartile range (25th to 75th percentiles), and the minimum and maximum non-outlier values (whiskers). Outliers, defined as values exceeding 1.5 times the interquartile range, are not shown. Only units that responded significantly according to the ZETA test are included. Numbers on top indicate the fraction of ZETA-significant units per category. **b,** Statistical assessment of the difference in response onset time between response categories. The matrix shows p-values derived from a mixed-effect regression model (Methods). P-values are visualized on a log10 color scale and are to be interpreted from column to row (red: significantly increased latency, blue: significantly decreased, grey: no significant difference). Only data from responsive, i.e. ZETA significant, units, is considered. **c,** Criteria for determining a suitable number of dataROI clusters to partition the PFC flatmap into modules based on the dataROI’s response category enrichment profiles (Fig. 5f,g). Vertical red dashed line marks the four clusters (modules) used. *Left*, Euclidean distance in metric space between the last pair of clusters joined as a function of the number of clusters (modules) defined when hierarchically clustering the enrichment profiles of dataROI’s nodes (read graph from right to left). Lower values indicate more homogenous clusters^47^. *Right*, Dunn index as a function of the number of clusters. The Dunn index is the ratio between minimal between-cluster distance and maximal with-cluster distance, with a higher Dunn index implying more compact and well-separated clusters ^48^. **d,** Copy of Supplementary Data Fig. 8a, data availability for the GaoROIs parcellating the PFC flatmap. GaoROIs with both hierarchy score and category enrichment profile are colored according to the PFC subregion where most of the units were recorded. White GaoROIs: no hierarchy score + too low (n < 20) unit count; black dots: GaoROI with hierarchy score but too low (n < 20) unit count; black stars: GaoROIs with category enrichment profile but lacking hierarchy score. **e to k,** Enrichment in response category vs PFC hierarchy score for ww categories 1–3 and 5–8. Pearson correlation: response category 1: R = 0.22; response category 2: R = 0.09; response category 3: R = -0.14; response category 5: R = -0.13; response category 6: R = -0.43; response category 7: R = -0.44; response category 8: R = 0.30. *Data:* dataset KI, PFC ww units, all brain (sub)regions and layers (n = 15,352 units); **a** and **b**. dataset KI, PFC ww units, restricted to deep layers (L5–6) for cortical regions (n = 12,373 units); **c** dataset KI, PFC ww units, deep layers, dataset KI (n = 9,232 units); **d** PFC hierarchy scores from Gao et al.^10^.

**Supplementary Data Fig. 10.**
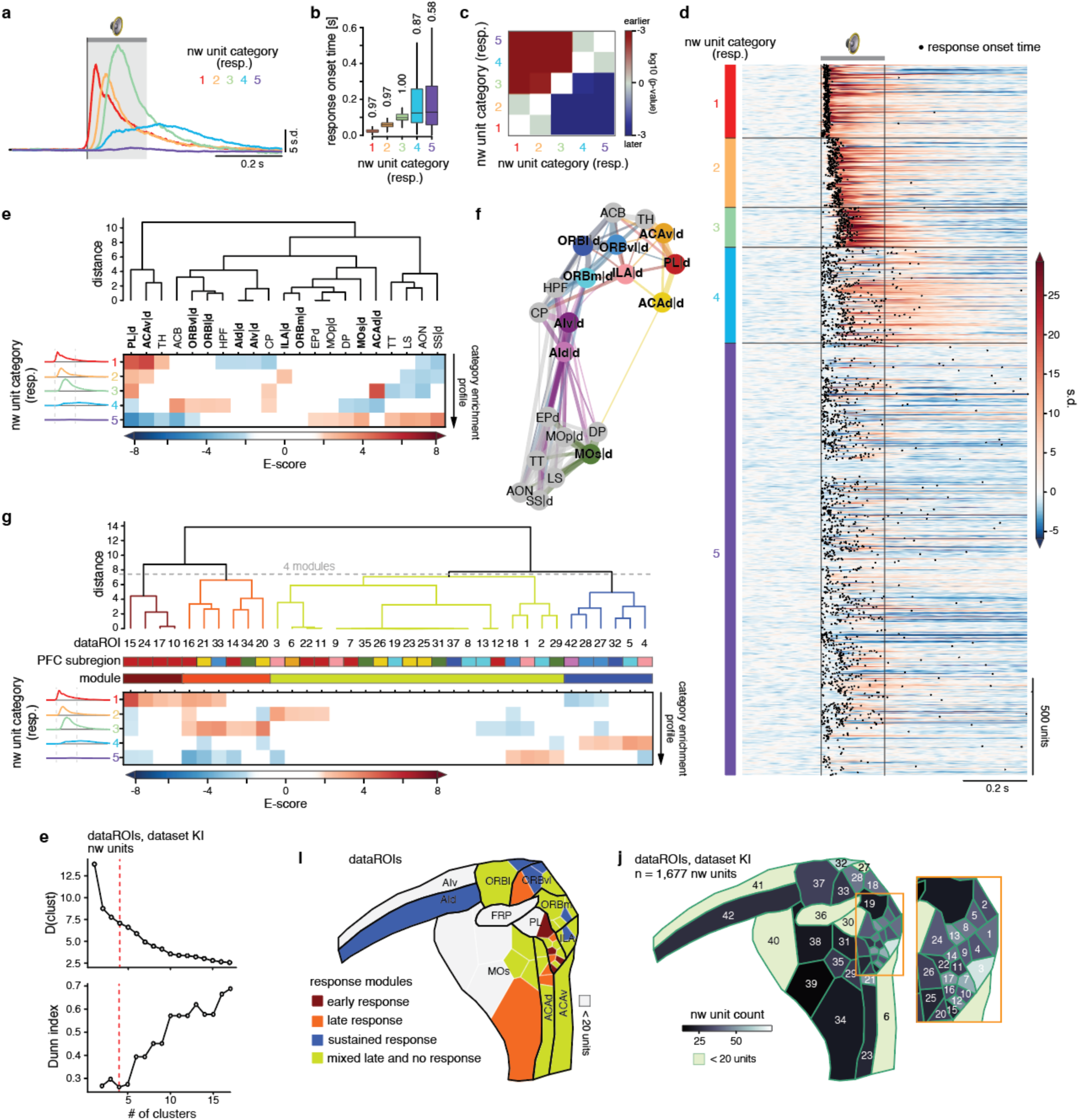
Enrichment in auditory response categories for nw units. **a,** The five response categories of nw units. Each colored line represents the normalized PSTH averaged across the units assigned to a response category. **b,** Response onset time (ZETA test,^38^) per nw response category. The whisker plot displays the median (center line), interquartile range (25th to 75th percentiles), and the minimum and maximum non-outlier values (whiskers). Outliers, defined as values exceeding 1.5 times the interquartile range, are not shown. Only units that responded significantly according to the ZETA test are included. Numbers on top indicate the fraction of ZETA-significant units per category. **c,** Statistical assessment of the difference in response latency between nw response categories. The matrix shows p-values derived from a mixed-effect regression model (Methods). P-values are visualized on a log10 color scale and are to be interpreted from column to row (red: significantly increased latency, blue: significantly decreased, grey: no significant difference). Only data from responsive, i.e. ZETA significant, units, is considered. **d,** Normalized PSTHs of single nw units arranged according to response category (1–5). Gray horizontal bar: tone presentation (200 ms); black vertical lines: tone onset/offset. Black dots: the time point of response peak onset (ZETA test^38^, significant units only, *p* < 0.01). **e,** *Bottom,* Response category enrichment profiles of brain (sub)regions for nw units. Bold: PFC subregions. *Top,* hierarchical tree (ward) derived from the enrichment profiles. *Left,* average normalized PSTH per category as in **a**. Dashed vertical lines: tone onset/offset. **f,** Graph display of the data in **e**. Nodes representing brain (sub)regions are arranged according to their first and second UMAP dimension, line width scales with cosine similarity between category enrichment profiles of brain regions (only shown for similarities > 0.1). Bold: PFC subregions. **g,** Clustering of PFC dataROIs (Fig. 4b) into response modules based on their response category enrichment profiles. *Top to bottom*, hierarchical tree (ward) derived from dataROI enrichment profiles; dataROI ID numbers; PFC subregion identity of the dataROIs (colored as in Fig. 4b); response modules; response category enrichment profiles of the PFC’s dataROIs. **h,** Criteria for determining a suitable number of dataROI clusters to partition the PFC flatmap into modules based on the dataROI’s response category enrichment profiles (**g**). Vertical red dashed line marks the four clusters (modules) used. *Top*, Euclidean distance in metric space between the last pair of clusters joined as a function of the number of clusters (modules) defined when hierarchically clustering the enrichment profiles of dataROI’s nodes. Lower values indicate more homogenous clusters^47^. *Bottom*, Dunn index as a function of the number of clusters. A higher Dunn index implies more compact and well-separated clusters^48^. **i,** PFC flatmap with dataROIs (white outlines) colored according to response module. Black outlines: cytoarchitecturally defined PFC subregions. **j,** Flatmap of the PFC with the count per dataROI of nw units (deep layers, L5–6) in dataset KI. Each dataROI (green outlines) is identified by an ID number. Box: enlargement of the corresponding frame on the flatmap. *Data:* dataset KI, nw units, all brain (sub)regions and layers, n = 3,653; **a**–**d** dataset KI, nw units, all brain (sub)regions, for cortex restricted to deep layers (L5–6), n = 1,486 units; **e** and dataset KI, nw units, PFC (sub)regions, restricted to deep layers (L5–6), n = 1,677 units; **g** to **j**

**Supplementary Data Table 1.** Enrichment p-values for enrichment matrix in Supplementary Data Fig. 3d P-values for each enrichment value in Supplementary Data Fig. 3d.

|  | spontaneous unit category |  |  |  |  |  |  |  |
| --- | --- | --- | --- | --- | --- | --- | --- | --- |
| Layer | 1 | 2 | 3 | 4 | 5 | 6 | 7 | 8 |
| 23 | 0,0265 | 0,4315 | 0,001 | 0 | 0 | 0,0045 | 0 | 0 |
| 4 | 0,125 | 0,1085 | 0,328 | 0,046 | 0,409 | 0,395 | 0,09 | 0,1915 |
| 1 | 0 | 0,008 | 0,0775 | 0,496 | 0,3185 | 0,3485 | 0,29 | 0,0225 |
| 5 | 0 | 0 | 0,1135 | 0,201 | 0,1305 | 0 | 0 | 0,045 |
| 6 | 0 | 0 | 0 | 0 | 0,019 | 0,211 | 0 | 0 |

**Supplementary Data Table 2.**
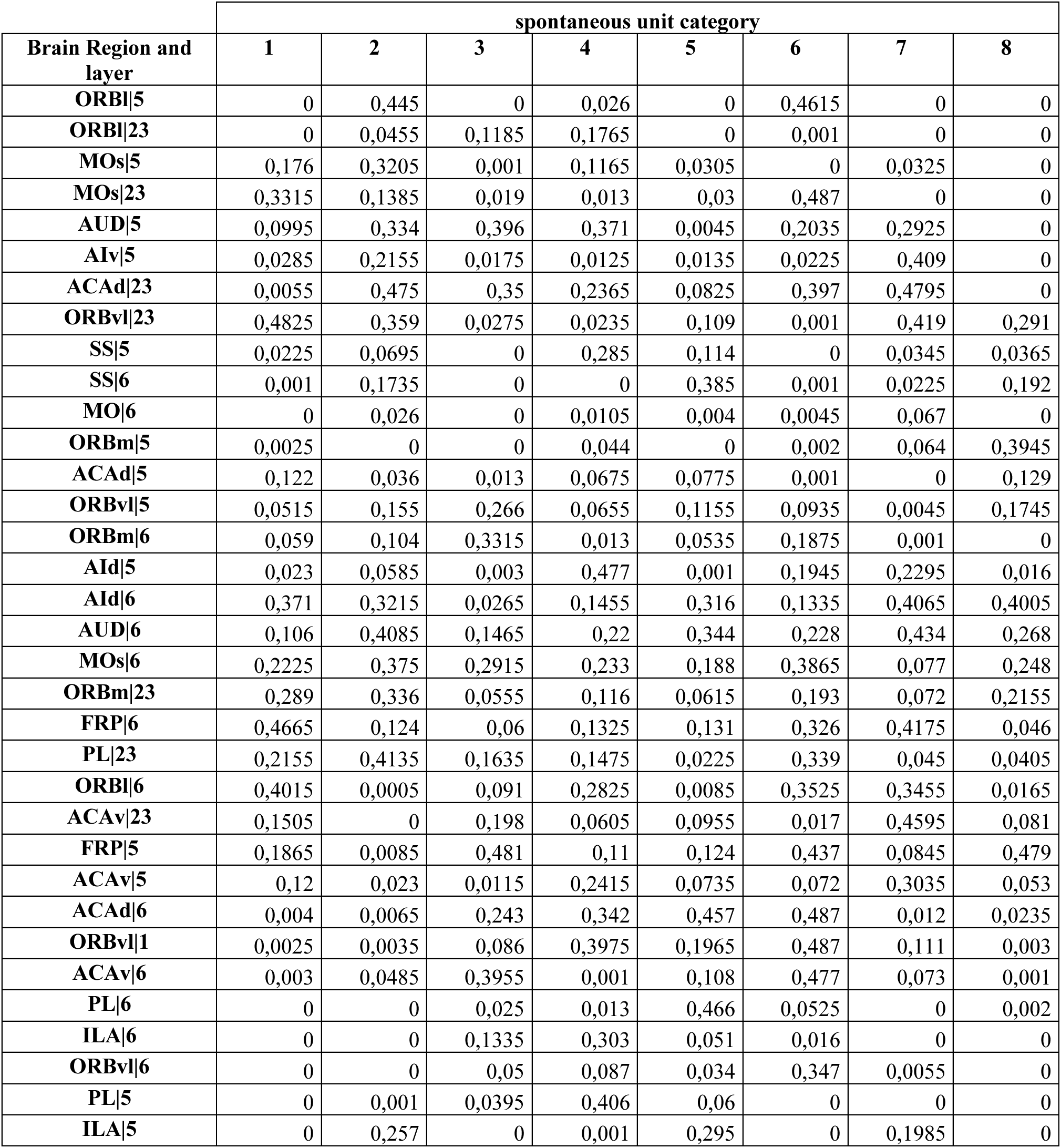
Enrichment p-values for enrichment matrix in Supplementary Data Fig. 3e P-values for each enrichment value in Supplementary Data Fig. 3e.

**Supplementary Data Table 3.** Enrichment p-values for enrichment matrix in Fig. 2j P-values for each enrichment value in Fig. 2j.

| brain region | spontaneous unit category |  |  |  |  |  |  |  |
| --- | --- | --- | --- | --- | --- | --- | --- | --- |
|  | 1 | 2 | 3 | 4 | 5 | 6 | 7 | 8 |
| LS | 0 | 0 | 0 | 0,0645 | 0 | 0 | 0 | 0 |
| ORBm\|d | 0,1485 | 0,043 | 0,001 | 0,2795 | 0 | 0,0015 | 0 | 0,0095 |
| ILA\|d | 0 | 0,016 | 0 | 0,0545 | 0,192 | 0 | 0,001 | 0 |
| ORBvl\|d | 0 | 0 | 0,0625 | 0,0175 | 0,015 | 0,0995 | 0 | 0 |
| PL\|d | 0 | 0 | 0,4535 | 0,0395 | 0,114 | 0 | 0 | 0 |
| ACAd\|d | 0,003 | 0,001 | 0,0135 | 0,0765 | 0,1065 | 0,007 | 0 | 0,434 |
| EPd | 0,093 | 0,076 | 0,096 | 0,0015 | 0,355 | 0,015 | 0,0065 | 0,1635 |
| FRP\|d | 0,3675 | 0,2405 | 0,086 | 0,0495 | 0,047 | 0,3865 | 0,1465 | 0,109 |
| AId\|d | 0,0775 | 0,0645 | 0,001 | 0,308 | 0,0025 | 0,114 | 0,2755 | 0,038 |
| DP | 0,0125 | 0,4875 | 0,039 | 0,0315 | 0,3525 | 0,0135 | 0,1695 | 0 |
| ACAv\|d | 0,007 | 0,0055 | 0,059 | 0,024 | 0,022 | 0,13 | 0,091 | 0 |
| SS\|d | 0,001 | 0,4495 | 0 | 0 | 0,193 | 0 | 0,003 | 0,0505 |
| TH | 0,0025 | 0,01 | 0 | 0 | 0,187 | 0 | 0,063 | 0,2645 |
| MO\|d | 0 | 0,0155 | 0 | 0,0275 | 0,007 | 0,009 | 0,227 | 0 |
| MOs\|d | 0,1015 | 0,3995 | 0,002 | 0,066 | 0,1205 | 0 | 0,161 | 0 |
| AUD\|d | 0,4715 | 0,338 | 0,296 | 0,183 | 0,019 | 0,104 | 0,2665 | 0,0055 |
| AIv\|d | 0,0255 | 0,2405 | 0,0255 | 0,0305 | 0,014 | 0,029 | 0,401 | 0 |
| CP | 0 | 0,0475 | 0,0065 | 0,112 | 0 | 0,035 | 0,314 | 0 |
| ACB | 0 | 0,1775 | 0,13 | 0,047 | 0,008 | 0,139 | 0,4905 | 0 |
| HPF | 0 | 0,003 | 0,236 | 0,278 | 0,1325 | 0,4415 | 0,0555 | 0,005 |
| ORBl\|d | 0 | 0,0615 | 0,001 | 0,0915 | 0 | 0,427 | 0,0005 | 0 |
| PIR | 0,005 | 0,106 | 0,002 | 0,153 | 0,0085 | 0,019 | 0 | 0 |
| AON | 0 | 0 | 0,4355 | 0,4615 | 0 | 0 | 0 | 0,015 |
| TT | 0 | 0,0715 | 0 | 0 | 0,003 | 0,1195 | 0 | 0,0195 |

**Supplementary Data Table 4.**
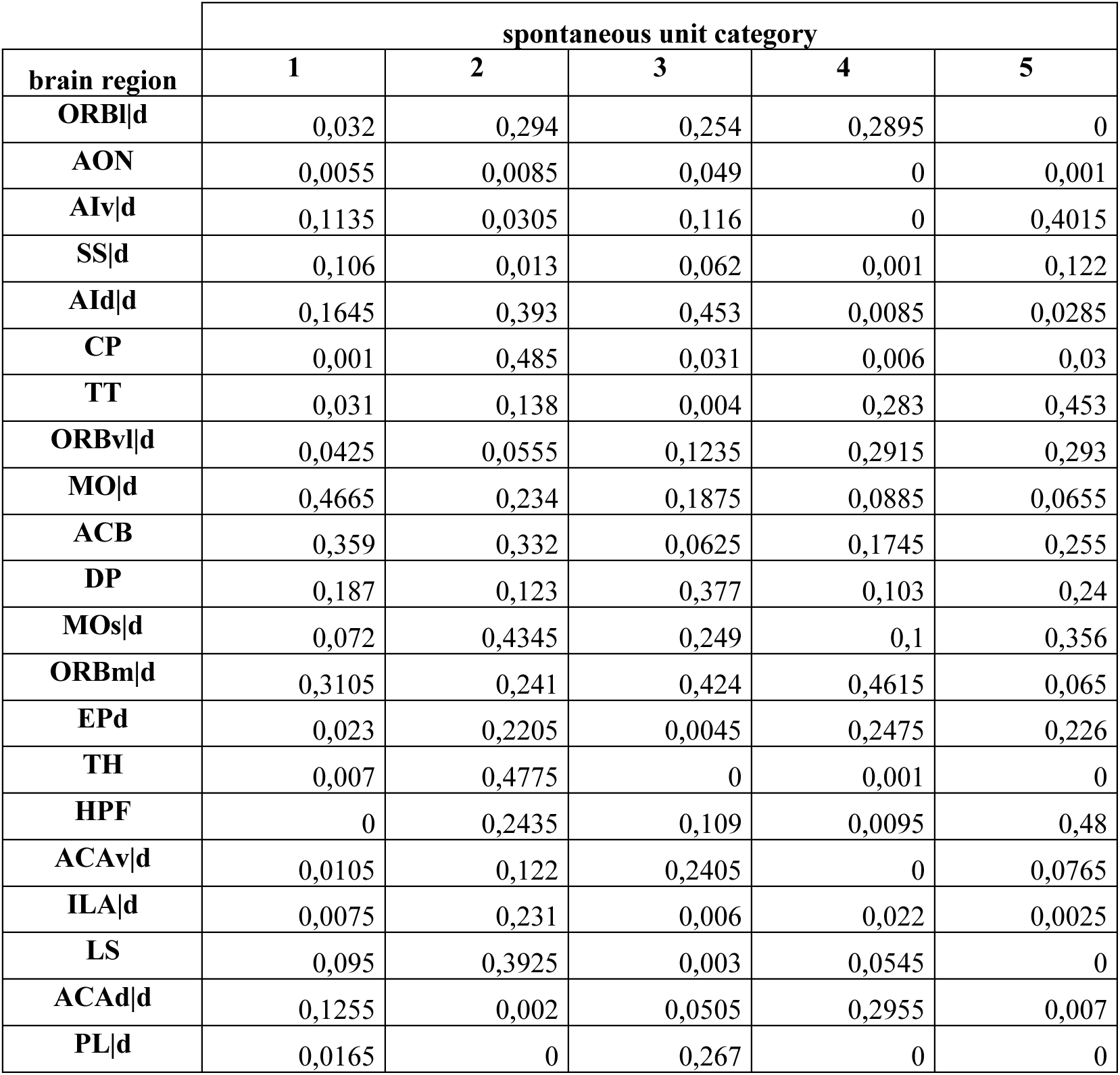
Enrichment p-values for enrichment matrix in Supplementary Data Fig. 5g P-values for each enrichment value in Supplementary Data Fig. 5g.

**Supplementary Data Table 5.** No correlation between enrichment and cortical hierarchy when restricting to PFC subregions Pearsons’ correlation coefficient (R) and p-values.

| dataset KI |  |  |  |  |  |  |  |  |  |  |  |  |  |  |
| --- | --- | --- | --- | --- | --- | --- | --- | --- | --- | --- | --- | --- | --- | --- |
|  | ww spontaneous unit category |  |  |  |  |  |  |  | nw spontaneous unit category |  |  |  |  | h-score |
| PFC subregion | 1 | 2 | 3 | 4 | 5 | 6 | 7 | 8 | 1 | 2 | 3 | 4 | 5 |  |
| ACAd | 3,36 | 2,91 | -1,50 | 0,62 | 1,77 | -2,75 | -4,47 | -0,97 | 1,16 | 3,35 | -1,77 | -1,01 | -2,47 | 0,36 |
| ACAv | 2,62 | 2,53 | 2,37 | 2,02 | -2,29 | -1,29 | -1,63 | -3,30 | 2,67 | 0,92 | 0,69 | -3,39 | -1,17 | 0,25 |
| AId | 1,01 | 1,08 | -2,26 | -0,17 | 2,21 | 1,08 | -0,35 | -1,28 | -0,97 | 0,00 | -0,35 | 2,51 | -1,50 | 0,13 |
| AIv | -2,04 | -1,01 | 2,96 | -1,99 | -2,67 | -1,77 | -0,11 | 3,82 | -1,11 | -1,94 | -1,25 | 4,11 | 0,48 | 0,33 |
| ILA | 8,72 | 1,79 | 6,17 | -2,05 | 0,45 | -3,51 | -2,73 | -9,12 | 2,26 | -0,26 | 2,52 | -2,10 | -2,98 | 0,23 |
| MOs | 2,27 | 0,76 | -3,29 | -0,98 | -0,23 | -3,42 | -0,28 | 3,71 | -1,31 | 0,04 | 0,79 | 0,87 | -0,63 | 0,32 |
| ORBI | -3,66 | 1,95 | -4,11 | 1,94 | -3,92 | 0,94 | 2,04 | 6,43 | -1,06 | 0,24 | -1,09 | -0,86 | 3,05 | 0,20 |
| ORBm | -0,02 | -1,72 | 2,69 | -0,57 | 4,97 | -2,67 | -2,80 | -3,67 | -0,87 | -0,18 | -0,09 | -0,14 | 1,58 | 0,23 |
| ORBvl | 1,91 | 4,18 | -2,74 | 3,77 | 1,06 | -1,19 | -2,87 | -2,36 | -1,75 | 1,20 | -1,13 | 1,14 | 0,30 | 0,39 |
| PL | 14,40 | 5,16 | 0,50 | 0,80 | -0,23 | -3,81 | -5,33 | -9,12 | 1,91 | 4,24 | 0,57 | -3,21 | -4,78 | 0,22 |
| FRP | -0,79 | 0,86 | -1,84 | -1,02 | 1,14 | 0,21 | -0,95 | 1,96 | NaN | NaN | NaN | NaN | NaN | 0,23 |
| Pearson (R) | -0,12 | 0,13 | -0,11 | 0,20 | -0,06 | -0,37 | -0,23 | 0,22 | -0,19 | 0,09 | -0,39 | 0,21 | 0,08 |  |
| sample size | 11 | 11 | 11 | 11 | 11 | 11 | 11 | 11 | 10 | 10 | 10 | 10 | 10 |  |
| t score | -0,36 | 0,39 | -0,34 | 0,60 | -0,19 | -1,18 | -0,72 | 0,68 | -0,55 | 0,25 | -1,19 | 0,61 | 0,21 |  |
| p-value | 0,73 | 0,71 | 0,74 | 0,56 | 0,85 | 0,27 | 0,49 | 0,52 | 0,60 | 0,81 | 0,27 | 0,56 | 0,84 |  |

| dataset IBL |  |  |  |  |  |  |  |  |  |  |  |  |  |  |
| --- | --- | --- | --- | --- | --- | --- | --- | --- | --- | --- | --- | --- | --- | --- |
|  | ww spontaneous unit category |  |  |  |  |  |  |  | nw spontaneous unit category |  |  |  |  | h-score |
| PFC subregion | 1 | 2 | 3 | 4 | 5 | 6 | 7 | 8 | 1 | 2 | 3 | 4 | 5 |  |
| ACAd | 1,51 | 3,62 | 0,02 | 1,58 | 3,00 | -1,65 | -2,39 | -4,00 | -1,20 | 1,30 | -1,84 | 1,17 | 1,10 | 0,36 |
| ACAv | 5,48 | -0,25 | -1,86 | -1,79 | 4,72 | 0,68 | -2,03 | -3,25 | 0,60 | 0,75 | -1,65 | 0,24 | 0,12 | 0,25 |
| AIv | 2,21 | 0,60 | 3,85 | -1,29 | 1,10 | -1,21 | -1,00 | -3,72 | -2,72 | 0,91 | -0,10 | 1,19 | 1,25 | 0,33 |
| ILA | 13,58 | 3,24 | 0,10 | -1,87 | 2,13 | -2,26 | -2,94 | -6,00 | -0,37 | 1,23 | 0,71 | -0,96 | -1,01 | 0,23 |
| MOs | 2,19 | 4,29 | 2,68 | -3,01 | 1,04 | -3,57 | 1,18 | -3,03 | -2,73 | -0,96 | 0,86 | 0,19 | 3,71 | 0,32 |
| ORBm | 6,48 | 4,00 | -0,03 | -0,76 | 0,51 | -2,05 | -1,00 | -2,86 | 0,01 | 1,17 | -1,34 | -0,69 | 1,03 | 0,23 |
| ORBvl | 5,70 | 4,58 | 0,56 | -0,09 | -2,66 | -3,30 | -2,76 | 0,90 | -1,71 | 1,57 | 0,96 | 0,26 | -1,38 | 0,39 |
| PL | 6,93 | 0,27 | 0,39 | -0,43 | 1,41 | -2,57 | -1,26 | -3,01 | 1,30 | 1,02 | -0,46 | -1,68 | -0,72 | 0,22 |
| AId | 1,81 | 2,98 | 1,99 | -2,56 | -2,50 | -2,05 | -0,22 | 1,79 | NaN | NaN | NaN | NaN | NaN | 0,13 |
| ORBI | -0,16 | 1,57 | 0,94 | -0,64 | 1,83 | -1,69 | -1,33 | -1,21 | NaN | NaN | NaN | NaN | NaN | 0,20 |
| Pearson (R) | -0,11 | 0,29 | 0,13 | 0,44 | 0,05 | -0,19 | -0,24 | -0,19 | -0,77 | 0,01 | 0,23 | 0,80 | 0,19 |  |
| sample size | 10 | 10 | 10 | 10 | 10 | 10 | 10 | 10 | 8 | 8 | 8 | 8 | 8 |  |
| t score | -0,30 | 0,86 | 0,36 | 1,40 | 0,14 | -0,55 | -0,70 | -0,54 | -2,93 | 0,02 | 0,57 | 3,26 | 0,46 |  |
| p-value | 0,77 | 0,42 | 0,73 | 0,20 | 0,89 | 0,60 | 0,50 | 0,60 | 0,03 | 0,99 | 0,59 | 0,02 | 0,66 |  |

**Supplementary Data Table 6.** Enrichment p-values for enrichment matrix in Fig. 4c P-values for each enrichment value in Fig. 4c.

| dataROI<br>ID | spontaneous unit category |  |  |  |  |  |  |  |
| --- | --- | --- | --- | --- | --- | --- | --- | --- |
|  | 1 | 2 | 3 | 4 | 5 | 6 | 7 | 8 |
| 40\|d | 0 | 0,0015 | 0 | 0,0795 | 0,043 | 0,3095 | 0,0515 | 0 |
| 32\|d | 0 | 0,4265 | 0 | 0,169 | 0 | 0,123 | 0 | 0 |
| 41\|d | 0 | 0,1745 | 0,0115 | 0,0025 | 0,0115 | 0,1295 | 0,3315 | 0 |
| 35\|d | 0,0255 | 0,057 | 0,397 | 0,4725 | 0,357 | 0,219 | 0,217 | 0 |
| 34\|d | 0,001 | 0,294 | 0,045 | 0,1945 | 0,4015 | 0,4175 | 0,3415 | 0 |
| 23\|d | 0,001 | 0,4295 | 0,011 | 0,275 | 0,0065 | 0,2155 | 0,241 | 0,001 |
| 42\|d | 0,2725 | 0,423 | 0 | 0,055 | 0,0065 | 0,0045 | 0,2995 | 0,22 |
| 18\|d | 0,406 | 0,09 | 0,079 | 0,0815 | 0,0125 | 0,004 | 0,4915 | 0 |
| 16\|d | 0,081 | 0,316 | 0,1965 | 0,2695 | 0,0135 | 0,2855 | 0,0075 | 0 |
| 5\|d | 0,05 | 0,0035 | 0,0075 | 0,041 | 0 | 0,003 | 0,126 | 0,059 |
| 2\|d | 0,0025 | 0,0765 | 0,2515 | 0,055 | 0,001 | 0,365 | 0,206 | 0,0515 |
| 36\|d | 0,0305 | 0,068 | 0,3225 | 0,4465 | 0,075 | 0,4135 | 0,0965 | 0,4995 |
| 21\|d | 0,0195 | 0,031 | 0,484 | 0,119 | 0,1415 | 0,242 | 0,067 | 0,2585 |
| 38\|d | 0,065 | 0,0265 | 0,2685 | 0,003 | 0,2245 | 0,477 | 0,384 | 0,267 |
| 30\|d | 0,012 | 0,0325 | 0,0795 | 0 | 0,1345 | 0,335 | 0,3725 | 0,011 |
| 39\|d | 0,105 | 0,0235 | 0,142 | 0,055 | 0,292 | 0,3325 | 0 | 0,21 |
| 22\|d | 0,412 | 0,0035 | 0,2525 | 0,302 | 0,178 | 0,1405 | 0,105 | 0,082 |
| 28\|d | 0,025 | 0 | 0,151 | 0,136 | 0,0695 | 0,0595 | 0,2085 | 0,1215 |
| 15\|d | 0,0155 | 0,1 | 0,353 | 0,3655 | 0,178 | 0,1075 | 0,1 | 0,437 |
| 24\|d | 0,042 | 0,2905 | 0,0675 | 0,2505 | 0,238 | 0,4795 | 0,494 | 0,242 |
| 31\|d | 0,236 | 0,2145 | 0,3655 | 0,2865 | 0,1915 | 0,124 | 0,0375 | 0,3095 |
| 20\|d | 0,395 | 0,094 | 0,1885 | 0,2705 | 0,2855 | 0,478 | 0,42 | 0,4075 |
| 14\|d | 0,2975 | 0,072 | 0,271 | 0,0955 | 0,286 | 0,1625 | 0,1305 | 0,1275 |
| 27\|d | 0,2745 | 0,4315 | 0,0555 | 0,463 | 0,031 | 0,0885 | 0,2625 | 0,421 |
| 29\|d | 0,2745 | 0,1715 | 0,22 | 0,1635 | 0,3315 | 0,0605 | 0,0455 | 0,0875 |
| 25\|d | 0,2955 | 0,0775 | 0,3735 | 0,3565 | 0,43 | 0,347 | 0,027 | 0,1265 |
| 33\|d | 0,214 | 0,0505 | 0,2555 | 0,0155 | 0,4265 | 0,339 | 0,008 | 0,2955 |
| 19\|d | 0,0935 | 0,273 | 0,2795 | 0,0235 | 0,05 | 0,0285 | 0,0705 | 0,141 |
| 6\|d | 0,4525 | 0,13 | 0,091 | 0,018 | 0,047 | 0,261 | 0,441 | 0,0235 |
| 13\|d | 0,2355 | 0,406 | 0,024 | 0,346 | 0,012 | 0,1395 | 0,3185 | 0,1295 |
| 37\|d | 0,069 | 0,064 | 0,477 | 0,175 | 0,0065 | 0,2485 | 0,2555 | 0,001 |
| 26\|d | 0 | 0,3085 | 0,0285 | 0,247 | 0,0635 | 0,0375 | 0 | 0,1 |
| 10\|d | 0 | 0,4435 | 0,0575 | 0,223 | 0,263 | 0,471 | 0,0625 | 0 |
| 11\|d | 0,003 | 0,112 | 0,2715 | 0,1185 | 0,3255 | 0,26 | 0,0095 | 0 |
| 7\|d | 0 | 0,3905 | 0,065 | 0,2425 | 0,21 | 0,135 | 0,001 | 0 |
| 17\|d | 0 | 0,0505 | 0,0165 | 0,0225 | 0,163 | 0,1255 | 0,035 | 0 |
| 12\|d | 0,0105 | 0,3025 | 0,0335 | 0,11 | 0,201 | 0,0385 | 0,254 | 0,002 |
| 9\|d | 0,0195 | 0,26 | 0,0275 | 0,422 | 0,135 | 0,1505 | 0,4285 | 0,0025 |
| 8\|d | 0,3985 | 0,1555 | 0,001 | 0,0465 | 0,0915 | 0,13 | 0,1225 | 0,043 |
| 1\|d | 0,063 | 0,4675 | 0 | 0,381 | 0,423 | 0,452 | 0,4545 | 0 |
| 4\|d | 0 | 0,026 | 0 | 0,036 | 0,4385 | 0,123 | 0,384 | 0 |
| 3\|d | 0 | 0,4355 | 0,0095 | 0,0295 | 0,1705 | 0,411 | 0,1235 | 0 |

**Supplementary Data Table 7.** Enrichment p-values for enrichment matrix in Supplementary Data Fig. 7e P-values for each enrichment value in Supplementary Data Fig. 7e.

|  | spontaneous unit category |  |  |  |  |  |  |  |
| --- | --- | --- | --- | --- | --- | --- | --- | --- |
| GaoROI ID | 1 | 2 | 3 | 4 | 5 | 6 | 7 | 8 |
| 23\|d | 0 | 0,3245 | 0,001 | 0,22 | 0 | 0,056 | 0 | 0 |
| 10\|d | 0 | 0,0115 | 0,0205 | 0,4355 | 0,003 | 0,4475 | 0 | 0 |
| 55\|d | 0,127 | 0,076 | 0,0015 | 0,0775 | 0,02 | 0,055 | 0,0345 | 0,001 |
| 28\|d | 0 | 0,1895 | 0,001 | 0,136 | 0,1525 | 0,038 | 0,29 | 0 |
| 25\|d | 0,005 | 0,2805 | 0,353 | 0,3 | 0,087 | 0,2435 | 0,3625 | 0,0005 |
| 46\|d | 0,001 | 0,063 | 0,063 | 0,158 | 0,4785 | 0,2285 | 0,258 | 0 |
| 36\|d | 0,028 | 0,0055 | 0,41 | 0,1 | 0,3455 | 0,157 | 0,339 | 0,0395 |
| 38\|d | 0,0045 | 0,002 | 0,3535 | 0,001 | 0,315 | 0,2 | 0,161 | 0 |
| 32\|d | 0 | 0,0725 | 0,002 | 0,0105 | 0,0095 | 0,1725 | 0,493 | 0 |
| 48\|d | 0 | 0,078 | 0,236 | 0,2 | 0 | 0,32 | 0,1025 | 0 |
| 53\|d | 0,001 | 0,0235 | 0 | 0,0165 | 0,43 | 0,236 | 0,0385 | 0 |
| 58\|d | 0,004 | 0,0085 | 0 | 0,433 | 0,002 | 0,2025 | 0,194 | 0 |
| 33\|d | 0,484 | 0,047 | 0 | 0,1255 | 0,172 | 0,0745 | 0,395 | 0,047 |
| 37\|d | 0 | 0,3555 | 0,02 | 0,3605 | 0 | 0,0495 | 0,262 | 0,024 |
| 13\|d | 0 | 0,193 | 0,199 | 0,0915 | 0,01 | 0,0145 | 0,0735 | 0,3785 |
| 49\|d | 0,038 | 0,13 | 0,2465 | 0,248 | 0 | 0,4175 | 0,4025 | 0,0035 |
| 9\|d | 0,4805 | 0,1155 | 0,389 | 0,1885 | 0,0095 | 0,189 | 0,182 | 0,017 |
| 14\|d | 0,383 | 0,4405 | 0,0995 | 0,1755 | 0,001 | 0,3045 | 0,0055 | 0,032 |
| 7\|d | 0,191 | 0,098 | 0,1665 | 0,013 | 0,3505 | 0,4705 | 0 | 0,198 |
| 47\|d | 0,451 | 0,3575 | 0,1945 | 0,0045 | 0,1665 | 0,2655 | 0,0765 | 0,032 |
| 24\|d | 0,2305 | 0,202 | 0,465 | 0,0135 | 0,1655 | 0,4385 | 0,465 | 0,0035 |
| 12\|d | 0,3715 | 0,237 | 0,184 | 0,0185 | 0,3015 | 0,4565 | 0,2555 | 0,136 |
| 43\|d | 0,113 | 0,2955 | 0,387 | 0,002 | 0,038 | 0,3555 | 0,187 | 0,1945 |
| 5\|d | 0,4225 | 0,2445 | 0,4685 | 0,0215 | 0,332 | 0,349 | 0,0265 | 0,096 |
| 11\|d | 0,079 | 0,431 | 0,128 | 0,0795 | 0,3905 | 0,395 | 0,021 | 0,427 |
| 42\|d | 0,0855 | 0,23 | 0,16 | 0,2925 | 0,039 | 0,21 | 0,0845 | 0,196 |
| 60\|d | 0,2095 | 0,194 | 0,2215 | 0,283 | 0,071 | 0,3485 | 0,133 | 0,151 |
| 21\|d | 0,4785 | 0,1345 | 0,004 | 0,4815 | 0,077 | 0,497 | 0,1355 | 0,408 |
| 35\|d | 0,094 | 0,346 | 0,402 | 0,434 | 0,276 | 0,017 | 0,3105 | 0,3135 |
| 2\|d | 0,2835 | 0,144 | 0,343 | 0,1975 | 0,439 | 0,001 | 0,2915 | 0,0205 |
| 6\|d | 0,001 | 0,015 | 0,332 | 0,0765 | 0,1445 | 0,283 | 0,1495 | 0,1265 |
| 8\|d | 0,303 | 0,0075 | 0,414 | 0,33 | 0,367 | 0,3675 | 0,213 | 0,077 |
| 56\|d | 0,034 | 0 | 0,1625 | 0,3245 | 0,131 | 0,0695 | 0,0595 | 0,008 |
| 40\|d | 0 | 0,2345 | 0,2205 | 0,1875 | 0,439 | 0,4825 | 0,2735 | 0 |
| 19\|d | 0 | 0,343 | 0,262 | 0,0915 | 0,2175 | 0,1645 | 0,026 | 0 |
| 59\|d | 0 | 0,027 | 0,0525 | 0,276 | 0,007 | 0,4215 | 0,006 | 0 |
| 20\|d | 0 | 0,002 | 0,1565 | 0,055 | 0,2225 | 0,4955 | 0,0155 | 0 |
| 41\|d | 0 | 0,0055 | 0,4535 | 0,0585 | 0,4935 | 0,3365 | 0,033 | 0 |
| 34\|d | 0 | 0,064 | 0,001 | 0,0985 | 0,0365 | 0,1085 | 0,0505 | 0 |

## References

1. Fuster, J. M. The Prefrontal Cortex. (Elsevier, 2015).

2. Carlen, M. What constitutes the prefrontal cortex? Science 358, 478–482 (2017).

3. Wang, X. et al. Three-dimensional intact-tissue sequencing of single-cell transcriptional states. Science 361, eaat5691 (2018).

4. Ortiz, C. et al. Molecular atlas of the adult mouse brain. Sci Adv 6, eabb3446 (2020).

5. Bhattacherjee, A. et al. Spatial transcriptomics reveals the distinct organization of mouse prefrontal cortex and neuronal subtypes regulating chronic pain. Nat Neurosci 26, 1880–1893 (2023).

6. Brodmann, K. Vergleichende Lokalisationslehre der Grosshirnrinde in ihren Prinzipien dargestellt auf Grund des Zellenbaues / [K. Brodmann]. (1909).

7. Franklin, K. B. J. & Praxinos, G. The Mouse Brain in Stereotaxic Coordinates. (Academic Press, 2007).

8. Van De Werd, H. J. J. M. & Uylings, H. B. M. Comparison of (stereotactic) parcellations in mouse prefrontal cortex. Brain Struct Funct 219, 433–459 (2014).

9. Harris, J. A. et al. Hierarchical organization of cortical and thalamic connectivity. Nature 575, 195–202 (2019).

10. Gao, L. et al. Single-neuron projectome of mouse prefrontal cortex. Nat Neurosci 25, 515–529 (2022).

11. Gao, L. et al. Single-neuron analysis of dendrites and axons reveals the network organization in mouse prefrontal cortex. Nat Neurosci 26, 1111–1126 (2023).

12. Zingg, B. et al. Neural Networks of the Mouse Neocortex. Cell 156, 1096–1111 (2014).

13. Ährlund-Richter, S. et al. A whole-brain atlas of monosynaptic input targeting four different cell types in the medial prefrontal cortex of the mouse. Nat Neurosci 22, 657–668 (2019).

14. Le Merre, P., Ährlund-Richter, S. & Carlén, M. The mouse prefrontal cortex: Unity in diversity. Neuron 109, 1925–1944 (2021).

15. Wilson, C. R. E., Gaffan, D., Browning, P. G. F. & Baxter, M. G. Functional localization within the prefrontal cortex: missing the forest for the trees? Trends Neurosci. 33, 533–540 (2010).

16. Christensen, A. J., Ott, T. & Kepecs, A. Cognition and the single neuron: How cell types construct the dynamic computations of frontal cortex. Current Opinion in Neurobiology 77, 102630 (2022).

17. Swindale, N. V., Spacek, M. A., Krause, M. & Mitelut, C. Spontaneous activity in cortical neurons is stereotyped and non-Poisson. Cereb Cortex 33, 6508–6525 (2023).

18. Maimon, G. & Assad, J. A. Beyond Poisson: Increased Spike-Time Regularity Across Primate Parietal Cortex. Neuron 62, 426–440 (2009).

19. Wang, X.-J. Macroscopic gradients of synaptic excitation and inhibition in the neocortex. Nat Rev Neurosci 21, 169–178 (2020).

20. Mochizuki, Y. et al. Similarity in Neuronal Firing Regimes across Mammalian Species. J. Neurosci. 36, 5736–5747 (2016).

21. Tolossa, G. B., Schneider, A. M., Dyer, E. L. & Hengen, K. B. A conserved code for anatomy: Neurons throughout the brain embed robust signatures of their anatomical location into spike trains. bioRxiv 2024.07.11.603152 (2024) doi:10.1101/2024.07.11.603152.

22. Siegle, J. H. et al. Survey of spiking in the mouse visual system reveals functional hierarchy. Nature 592, 86–92 (2021).

23. Murray, J. D. et al. A hierarchy of intrinsic timescales across primate cortex. Nat Neurosci 17, 1661–1663 (2014).

24. Zeisler, Z. R., Love, M., Rutishauser, U., Stoll, F. M. & Rudebeck, P. H. Consistent hierarchies of single-neuron timescales in mice, macaques and humans. bioRxiv 2024.10.30.621133 (2024) doi:10.1101/2024.10.30.621133.

25. Fuster, J. M. The prefrontal cortex--an update: time is of the essence. Neuron 30, 319–333 (2001).

26. Vyazovskiy, V. V. et al. Local sleep in awake rats. Nature 472, 443–447 (2011).

27. Goh, K.-I. & Barabási, A.-L. Burstiness and memory in complex systems. EPL 81, 48002 (2008).

28. Petersen, P. C., Siegle, J. H., Steinmetz, N. A., Mahallati, S. & Buzsáki, G. CellExplorer: A framework for visualizing and characterizing single neurons. Neuron 109, 3594–3608.e2 (2021).

29. Senzai, Y., Fernandez-Ruiz, A. & Buzsáki, G. Layer-Specific Physiological Features and Interlaminar Interactions in the Primary Visual Cortex of the Mouse. Neuron 101, 500–513.e5 (2019).

30. Zhang, S. et al. Organization of long-range inputs and outputs of frontal cortex for top-down control. Nature Neuroscience 19, 1733–1742 (2016).

31. Chaudhuri, R., Knoblauch, K., Gariel, M.-A., Kennedy, H. & Wang, X.-J. A Large-Scale Circuit Mechanism for Hierarchical Dynamical Processing in the Primate Cortex. Neuron 88, 419–431 (2015).

32. Salaj, D. et al. Spike frequency adaptation supports network computations on temporally dispersed information. eLife 10, e65459 (2021).

33. Dembrow, N. C., Zemelman, B. V. & Johnston, D. Temporal Dynamics of L5 Dendrites in Medial Prefrontal Cortex Regulate Integration Versus Coincidence Detection of Afferent Inputs. J. Neurosci. 35, 4501–4514 (2015).

34. Dégenètais, E., Thierry, A.-M., Glowinski, J. & Gioanni, Y. Electrophysiological Properties of Pyramidal Neurons in the Rat Prefrontal Cortex: An In Vivo Intracellular Recording Study. Cerebral Cortex 12, 1–16 (2002).

35. Harris, K. D. & Shepherd, G. M. G. The neocortical circuit: themes and variations. Nature Neuroscience 18, 170–181 (2015).

36. Moberg, S. & Takahashi, N. Neocortical layer 5 subclasses: From cellular properties to roles in behavior. Front. Synaptic Neurosci. 14, (2022).

37. Hafizi, H., et al. Inhibition-Dominated Rich-Club Shapes Dynamics in Cortical Microcircuits. *bioRxiv* 2021.05.07.443074 (2022) doi:10.1101/2021.05.07.443074.

38. Kupferschmidt, D. A. et al. Prefrontal Interneurons: Populations, Pathways, and Plasticity Supporting Typical and Disordered Cognition in Rodent Models. J. Neurosci. 42, 8468–8476 (2022).

39. Montijn, J. S. et al. A parameter-free statistical test for neuronal responsiveness. eLife 10, e71969 (2021).

## Additional references

40. Schomburg, E. W. et al. Theta Phase Segregation of Input-Specific Gamma Patterns in Entorhinal-Hippocampal Networks. Neuron 84, 470–485 (2014).

41. Osanai, H., Yamamoto, J. & Kitamura, T. Extracting electromyographic signals from multi-channel LFPs using independent component analysis without direct muscular recording. Cell Reports Methods 3, 100482 (2023).

42. International Brain Laboratory et al. A Brain-Wide Map of Neural Activity during Complex Behaviour. *bioRxiv* 2023.07.04.547681 (2023) doi:10.1101/2023.07.04.547681.

43. Shinomoto, S. et al. Relating Neuronal Firing Patterns to Functional Differentiation of Cerebral Cortex. PLOS Computational Biology 5, e1000433 (2009).

44. Kohonen, T. Essentials of the self-organizing map. Neural Networks 37, 52–65 (2013).

45. Kohonen, T. Self-Organizing Maps. vol. 30 (Springer, Berlin, Heidelberg, 2001).

46. Kind, M. C. & Brunner, R. J. SOMz: photometric redshift PDFs with self organizing maps and random atlas. Preprint at 10.48550/arXiv.1312.5753 (2013).

47. Ward Jr., J. H. Hierarchical Grouping to Optimize an Objective Function. Journal of the American Statistical Association 58, 236–244 (1963).

48. Thorndike, R. L. Who belongs in the family? Psychometrika 18, 267–276 (1953).

49. Dunn, J. C. Well-Separated Clusters and Optimal Fuzzy Partitions. Journal of Cybernetics 4, 95– 104 (1974).

50. Levy-Kramer, J. k-means-constrained. (2018).

51. Gillies, S. Shapely: manipulation and analysis of geometric objects. (2007).

52. Yu, Z. et al. Beyond t test and ANOVA: applications of mixed-effects models for more rigorous statistical analysis in neuroscience research. Neuron 110, 21–35 (2022).

53. Kuznetsova, A., Brockhoff, P. B. & Christensen, R. H. B. lmerTest Package: Tests in Linear Mixed Effects Models. Journal of Statistical Software 82, 1–26 (2017).

54. Halekoh, U. & Højsgaard, S. A Kenward-Roger Approximation and Parametric Bootstrap Methods for Tests in Linear Mixed Models – The R Package pbkrtest. Journal of Statistical Software 59, 1–32 (2014).

55. Model Selection and Multimodel Inference. (Springer, New York, NY, 2004). doi:10.1007/b97636.

